# Unsupervised reference-free inference reveals unrecognized regulated transcriptomic complexity in human single cells

**DOI:** 10.1101/2022.12.06.519414

**Authors:** Roozbeh Dehghannasiri, George Henderson, Rob Bierman, Tavor Baharav, Kaitlin Chaung, Peter Wang, Julia Salzman

**Affiliations:** Department of Biomedical Data Science, Stanford University, Stanford, CA 94305i; Department of Biochemistry, Stanford University, Stanford, CA 94305; Department of Electrical Engineering, Stanford University, Stanford, CA 94305; Department of Statistics and of Biology (by Courtesy), Stanford University, Stanford, CA 94305

**Author notes:** These authors contributed equally to this work (Roozbeh Dehghannasiri, George Henderson).

## Abstract

Myriad mechanisms diversify the sequence content of eukaryotic transcripts at both the DNA and RNA levels, leading to profound functional consequences. Examples of this diversity include RNA splicing and V(D)J recombination. Currently, these mechanisms are detected using fragmented bioinformatic tools that require predefining a form of transcript diversification and rely on alignment to an incomplete reference genome, filtering out unaligned sequences, potentially crucial for novel discoveries. Here, we present SPLASH+, significantly advancing biological discovery possible with SPLASH, our recently introduced efficient, reference-free statistical approach. Integrating a micro-assembly and biological interpretation framework, SPLASH+ enables new discoveries including broad and novel examples of transcript diversification in single cells *de novo*, without the need for cell type metadata, which is impossible with current algorithms. Applied to 10,326 primary human single cells across 19 tissues profiled with SmartSeq2, SPLASH+ discovers a set of splicing and histone regulators with highly conserved intronic regions that are themselves subject to complex splicing regulation. Additionally, it reveals unreported transcript diversity in the heat shock protein *HSP90AA1*, as well as diversification in centromeric RNA expression, V(D)J recombination, RNA editing, and repeat expansion, all missed by existing methods. SPLASH+ is highly efficient, enabling the discovery of an unprecedented breadth of RNA regulation and diversification in single cells through a new automated paradigm of unbiased transcriptomic analysis.

## Introduction

The diversity of transcripts in eukaryotes results from mechanisms such as alternative splicing, RNA editing, and alternative 5’ and 3’ UTR use as well as genetic changes in single cells including insertion of mobile elements, repeat expansions, or segmental duplications. The genome’s potential to create more than 10^13^ genetic variants through V(D)J recombination (Schroeder 2006) in the adaptive immune system is crucial for determining the specificity and efficacy of defense mechanisms against pathogens. Transcript diversification can have significant functional consequences, including causal links to various diseases from cancer to neurodegeneration (Kung, Maggi, and Weber 2018; Yum, Wang, and Kalsotra 2017; Bonnal, López-Oreja, and Valcárcel 2020; Ma et al. 2021)

Despite its importance to cell specialization, the extent to which transcript diversity is regulated in single cells remains a significant open question. Current computational approaches to detect transcript diversity in single cells are specialized for certain events such as alternative splicing (Olivieri, Dehghannasiri, and Salzman 2022; Buen Abad Najar et al. 2022) or V(D)J recombination reconstruction (Lindeman et al. 2018) and also rely heavily on references, beginning with the alignment of reads to a reference genome, thus censoring unmapped reads and introducing mapping biases. Other tasks, such as identifying somatically acquired repeats or RNA editing are not even attempted in scRNA-seq. Further, it is unclear whether these methods are sensitive to all events they aim to detect. For example, they may fail to align highly edited or spliced transcripts (Eisenberg and Levanon 2018), and most significantly, they are incapable of detecting sequences that are absent from a reference genome.

Statistics is at the core of inference for single-cell genomics. Yet, current genomic inference is conditional on the results of partially heuristic alignment algorithms, which often discard reads that do not map to the reference genome (Figure 1A). Moreover, these reference-first approaches might miss sequence diversity in underrepresented populations, as the current human reference genome is mostly European-derived (Sherman et al. 2019). The statistical tests downstream of alignment are typically parametric or require randomized resampling, leading to inaccurate or inefficient p-values. De novo assembly approaches (Cmero et al. 2021; Swanson et al. 2013), while attempting to create more accurate references, also introduce fundamental biases and unknown false positive and negative rates as shown by previous studies (Freedman, Clamp, and Sackton 2021). Even with a more accurate reference genome, alignment could still bias downstream statistical inference. Together, there is a strong argument to bypass reference alignment prior to statistical inference to find regulated sequence diversity through an unbiased and unified framework, changing the field’s “reference-first” paradigm to “statistics-first”.

**Figure 1.**
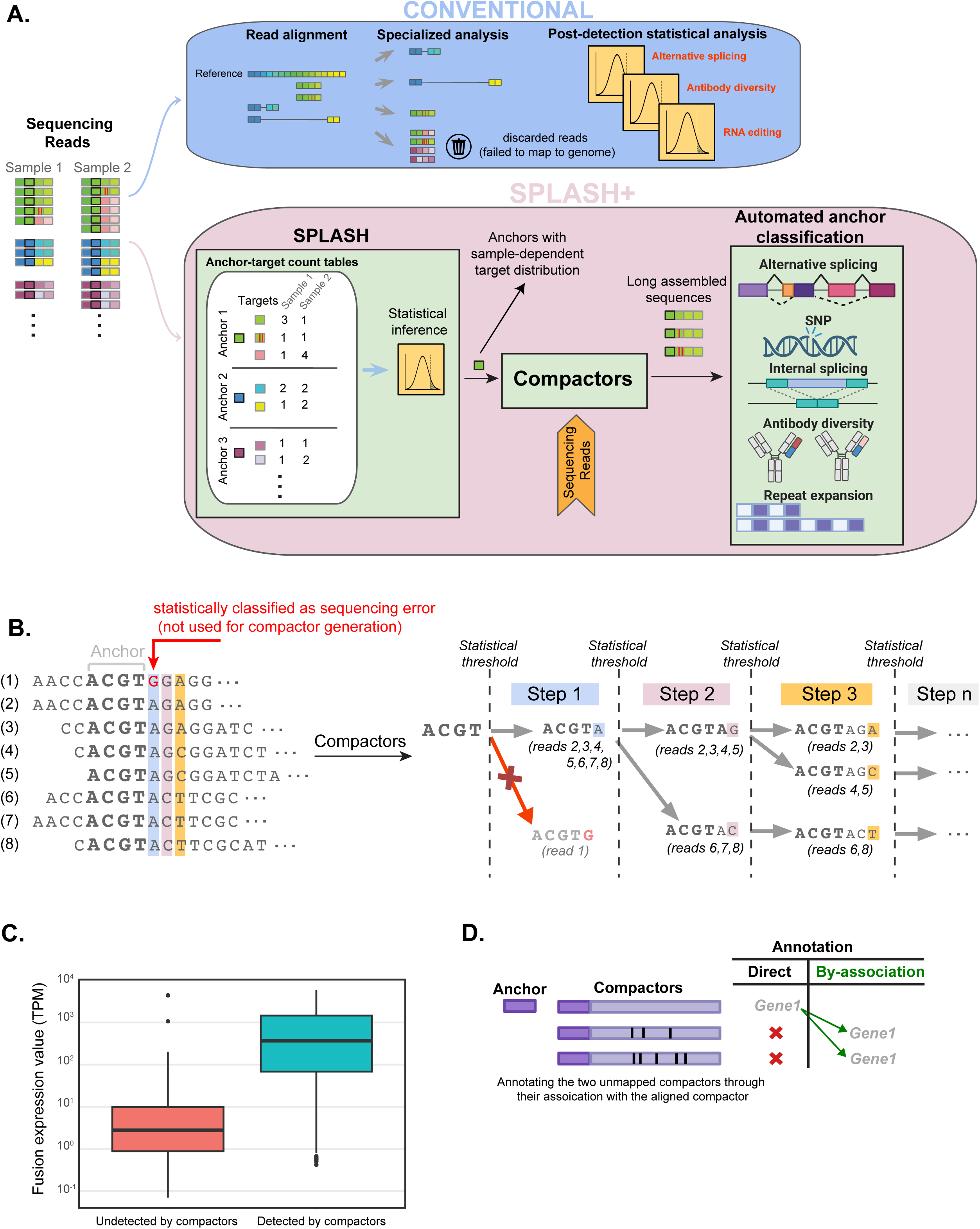
A unified reference-free approach to transcriptomic diversity analysis. (A) Current RNA-seq analysis methods, relying on read alignment to a reference genome, introduce biases and blindspots, and are tailored for specific applications. SPLASH offers a unified alignment-free solution for detecting myriad transcriptome diversifying mechanisms. SPLASH parses reads from all samples (cells) to extract constant kmers (anchors) that are followed by a set of diverse kmers (targets). For each anchor, SPLASH creates a contingency table and computes a closed-form p-value. Anchors with significant p-values are indicative of single-cell-dependent target distribution evidencing regulation and can subsequently be used for downstream analysis for a wide range of applications. SPLASH+ integrates SPLASH with a local assembly approach, called compactors, and an automated anchor classification based on compactors to facilitate biological interpretation and categorize anchors into biologically relevant events such as alternative splicing, SNP, and V(D)J recombination. (B) SPLASH-called anchors are extended to longer sequences or “compactors’’ through a statistically valid local assembly approach. The process involves collecting reads containing the anchor, recursively extending the sequence beyond the anchor by comparing nucleotide frequencies at each position against a threshold based on sequencing error, resulting in distinct branches that correspond to compactors for the anchor. (C) The boxplot showing the expression values (TPM) for the detected and undetected fusions by compactors in the fusion benchmarking dataset, suggesting that compactors provided evidence for the majority of sufficiently-expressed fusions. (D) Unmapped compactors can be *annotated by association* through other compactors linked to the same anchor.

SPLASH (Chaung et al. 2023) is a unified reference-free algorithm that performs statistical inference directly on raw sequencing reads to detect regulated sequence diversity. SPLASH’s core includes a statistical test to detect sample-specific sequence variation independent of any cell metadata such as cell type, by providing finite-sample valid p-value bounds, that unlike Pearson’s chi-squared test, better controls false positive calls under commonly used models such as negative binomial for scRNA-seq (Buen Abad Najar, Yosef, and Lareau 2020; Baharav, Tse, and Salzman 2023).

A weakness of SPLASH that limits its utility for targeted studies is the interpretation of its results. Sequence diversity can arise from a wide range of biological processes such as RNA splicing, RNA editing, repetitive region variation, and V(D)J recombination. As a result, SPLASH requires substantial manual effort to identify the cause of diversity and needs a systematic classification approach to categorize the detected sequence diversity into biologically meaningful events (e.g., RNA splicing) to draw meaningful biological insight. SPLASH also lacks sequence context as the standard output for SPLASH includes very short sequences (default k=27), which are insufficient for accurately attributing biological mechanisms that require longer sequences. This is especially problematic in identifying transcript isoforms involving alternative splicing of multiple exons or complex recombination events like V(D)J, requiring sufficiently long sequences to define variability of the Variable, Diversity, and Joining segments.

To address these shortcomings of SPLASH, in this manuscript, we build on SPLASH’s core to introduce SPLASH+, providing a systematic framework to analyze and interpret SPLASH’s output. This includes a new, reference-free statistical approach for *de novo* assembly of short sequences called by SPLASH to generate longer sequences called *compactors*, and also a framework to interpret and classify SPLASH’s results based on these compactor sequences (Figure 1A). We note that SPLASH+’s results are “reference-free” in the sense that statistical inference for detecting sample-specific sequence variation is completely independent of assembly and potential alignment biases. Alignment to the reference genome is used only after statistical inference in the subsequent classification step. Thus, SPLASH+ provides “the best of both worlds”: unbiased statistical guarantees of reference-free inference, along with the interpretability of reference-first approaches.

To systematically analyze transcript diversity in human cells, we applied SPLASH+ to 10,326 human cells across 346 cell types and 12 donors profiled with SmartSeq2 (SS2) from the *Tabula Sapiens* dataset (Tabula Sapiens Consortium* et al. 2022), a comprehensive human single-cell RNA sequencing dataset spanning multiple tissues and individuals. SPLASH+ reveals new insights into the regulation of transcript diversification in human single cells, including features of RNA splicing, RNA editing, and non-coding RNA expression missed by specialized, domain-specific bioinformatic pipelines. Novel findings include (i) regulated expression of repetitive loci such as *RNU6* variants and higher order repeats in centromeres; (ii) complex splicing programs, including un-annotated variants in genes like *CD47*, a major cancer immunotherapy target, and *RPS24*; (iii) pan-tissue regulation of splicing in splicing factors, histone modifications, and in the heat shock protein *HSP90AA1*; (iv) *de novo* rediscovery of immunoglobulin loci as the most transcriptionally diverse human loci with higher sensitivity than the current state-of-the-art for V(D)J rearrangement detection; and (v) the detection of single cells with transcribed repeat expansion and high levels of RNA editing. SPLASH+ makes unbiased discoveries by not relying on cell metadata or reference genomes. This is particularly important for scRNA-seq analysis, as cell types can be difficult to generate and remain imprecise (Zeng 2022) and can miss important variation within cell types, such as B cell receptor variation (Watson and Breden 2012). Our results demonstrate that SPLASH+ is a single algorithm that could replace numerous custom bioinformatic approaches for detecting different types of RNA variation and significantly expands our understanding of the transcriptome’s diversity.

### An integrated, reference free pipeline to discover regulated RNA expression

SPLASH (Chaung et al. 2023) is an unbiased reference-free algorithm that operates directly on raw sequencing data to identify differentially diversified sequences. This variation can be a signature of alternative RNA splicing, RNA editing, V(D)J recombination, or other mechanisms. SPLASH parses reads from all input samples and identifies specific *k*-mers (substrings of length *k*), called *anchors*, where each anchor is followed by a set of diverse k-mers, called *targets* (Methods, Figure 1A). SPLASH then performs a statistical test for each anchor under the null hypothesis that target count distribution is the same across all samples, providing a closed-form p-value bound for each anchor. Anchors with significant multiple testing-corrected p-values have sample-specific target expression, reflecting inter-cell expression variation.

Given that sequence diversity can arise from various mechanisms, SPLASH currently requires intensive manual interpretation to perform downstream analysis. Additionally, its output sequences (anchors and targets) are too short (both 27-mers by default) to provide sufficient context for inferring complex diversifying mechanisms (e.g., RNA alternative splicing involving multiple exons) or those spanning long loci (e.g., V(D)J rearrangement). These limitations hinder SPLASH’s potential to systematically analyze specific mechanisms (e.g., alternative splicing) in targeted studies.

To address these issues, we developed SPLASH+ by integrating SPLASH with a new seed-based statistical de novo assembly to establish longer sequence contexts, called *compactors,* for each short anchor called by SPLASH (Figure 1A). When compactors are built for each SPLASH-called anchor, they are used to classify the anchor into biologically-meaningful categories (Figure 1A). Integrating these two steps, SPLASH+ enhances the interpretability of SPLASH and facilitates targeted downstream analysis of specific events of biological interest.

To generate compactors, SPLASH-called anchors are used as seeds (Figure 1B; Methods). First, the reads containing each anchor are collected by parsing the input FASTQ files. Then, the anchor sequence is extended using a recursive branching rule, where each new branch at a given position corresponds to a specific nucleotide in the assembled sequence. Starting from the first position downstream of the anchor, we compute the frequency of each nucleotide in the given position across all collected reads for the anchor. A new branch is created for each nucleotide that has a frequency exceeding a certain threshold derived from probabilistic analysis (Methods), thereby minimizing the likelihood of generating a new branch due to sequencing error. For each created branch, reads containing the corresponding nucleotide are then propagated to that branch (Figure 1B). In the next iteration, nucleotide frequencies at the next position are evaluated per branch using the propagated reads, and the sequence is extended accordingly. This procedure continues recursively until a user-defined number of iterations is reached or the number of reads drops below a user-defined threshold. Upon completion, an assembled sequence, called a *compactor*, (along with the number of reads supporting it, as a measure of its abundance) is reported for each path of branches. Our statistical analysis providing a conservative closed-form null probability shows it is highly unlikely for a branch arising from sequencing error – i.e., sequencing-error-induced branches or false positive contigs (Methods).

While the main output of SPLASH includes anchor and target sequences, as a post-hoc annotation step, SPLASH (Chaung et al. 2023) can also build a single consensus sequence for each anchor in each sample by simply taking the plurality vote of the reads (Suppl. Figure 1A), i.e., each position in consensus sequence contains the nucleotide with the highest count in the sample. The consensus sequences are then mapped to the genome for gene name and alignment annotation. However, this approach has several drawbacks. A key issue with consensus is that it reports only one sequence per sample (cell), potentially failing to capture sequence diversity variation across samples. For example, the consensus sequence for an anchor might be identical across samples (incorrectly implying no sequence variation across samples), yet the contribution of each target can differ across samples (e.g., splicing between multiple isoforms or masking of mutations that have lower counts compared to the dominant variant) (Suppl. Figures 1B). Additionally, using a simple plurality vote across all reads can theoretically result in misassembly, where the represented variant does not exist in the sample (Suppl. Figure 1B). The new compactors approach in SPLASH+ addresses these issues by employing a statistical assembly approach that incorporates branching, allowing for multiple assembled sequences for each anchor per sample. Also, by considering only the subset of reads propagated to each branch, it avoids the misassembled sequences that can arise by the consensus approach. Importantly, compactors also preserve count information by reporting the number of reads supporting each compactor, which is crucial for subsequent classification of anchors to biologically relevant mechanisms such as RNA splicing or V(D)J recombination that requires considering the most abundant sequences. In contrast, the consensus approach in SPLASH loses this count information, resulting in a lack of automated, systematic anchor classification in SPLASH.

Compactors distinguish sequencing errors from inherent biological sequence variations, offering a straightforward and interpretable framework for dissecting transcriptome complexity and enabling the differentiation between different types of transcript diversification. To our knowledge, unlike any other de novo transcript assembly used for scRNA-seq, compactors can be statistically characterized to quantify the probability of generating an artifactual compactor due to sequencing error (Methods). Furthermore, compactors reduce the computational burden of downstream analysis, such as genome alignment. For example, in this study, compactors reduced the number of sequences 120 fold: from 183,471,175 raw reads to 1,515,555 compactors.

We tested if compactors could precisely reconstruct gene fusions using 5 simulated datasets from a benchmarking study (Haas et al. 2019). This widely-used dataset is a valuable resource for evaluating compactors, as it provides ground truth sequences (for gene fusions), which are essential for evaluating a sequence assembly method. We should note that there is no difference between gene fusions and other RNA variants (e.g., RNA splicing) from a sequence assembly viewpoint. We took the 27-mer immediately upstream of each fusion breakpoint as seeds and generated compactors (Suppl. Figure 2A, Methods). We then tested if generated compactors showed evidence for a fusion based on whether the compactor contained the fusion breakpoint (Suppl. Figure 2A). Compactors identified evidence for 57.8% (1,339 / 2,315) of total fusions (Suppl. Figure 2B; Suppl. Table 1). Compactor sensitivity surpassed that of two de novo assembly fusion detection methods (JAFFA-Assembly and TrinityFusion-D) in (Haas et al. 2019) and was comparable to the other two methods (TrinityFusion-C and TrinityFusion-UC) that exclusively utilized chimeric and unaligned reads, biased for fusion transcripts, and are highly computationally intensive (Figure S3 panel A in (Haas et al. 2019)). Our evaluation of compactors using a benchmarking dataset with ground truth sequences suggests that, despite being designed for general sequence assembly, compactors demonstrate comparable or even higher sensitivity for gene fusion detection than other de novo methods specifically optimized for this purpose. Notably, compactors detected the majority of sufficiently-expressed fusions as 98% of undetected fusions had TPM < 100 and only 2% of 965 fusions with TPM > 100 were undetected by compactors (Figure 1C).

After generating compactors for each SPLASH-called anchor, they are used to classify the anchor into one of 6 different categories (Methods, Suppl. Figure 3): splicing, internal splicing, base pair change, 3’UTR, centromere, and repeat. The classification of each anchor is based on the two compactors with the most aggregated assigned reads across all samples. This classification is either reference-free, utilizing string metrics such as Hamming and Levenshtein distances (for internal splicing and base pair change), or reference-based, by aligning to the human T2T reference genome using a spliced aligner such as STAR (Dobin et al. 2013) (for splicing, 3’UTR, centromere, and repeat).

An anchor is classified as splicing if STAR reports spliced alignment for at least one of the top two compactors (Methods). When both compactors lack splice junctions, the anchor is classified as internal splicing or base pair change, based on the string distances between the two compactors (Methods). The mapping positions of the remaining unclassified anchors are intersected with annotation databases for 3’ UTRs, centromeric repeats, and repeats to classify them accordingly. Through our classification, each compactor, even if it fails to map, can still be annotated by association with the annotation of the most abundant compactor for the anchor (*annotated-by-association*) (Figure 1D). This crucially enables the annotation of unaligned compactors for loci that are difficult to map, such as those involved in V(D)J recombination or noncoding RNAs with repetitive structures or multiple copies, such as spliceosomal snRNAs (as discussed later in the manuscript). As shown later in the manuscript, this can increase the sensitivity of SPLASH+ for detecting these variants. By performing reference alignment only after statistical inference for interpretability and automated anchor classification, SPLASH+ combines the best of two worlds: ensuring statistically valid and unbiased inference along with convenient interpretation and allows for a direct comparison of the improvements it offers over existing algorithms.

### SPLASH+ detects transcript diversity in repetitive RNA loci including centromeres and U6 spliceosomal RNA

We ran SPLASH+ on 10,326 cells profiled with SmartSeq2 from 19 tissues and 12 donors (29 donor-tissue pairs) and 346 cell types from the *Tabula Sapiens* Dataset (Tabula Sapiens Consortium* et al. 2022) (Suppl. Figure 4), including 10 tissues (e.g., blood, muscle, lung) with at least two donors, allowing us to analyze reproducibility, as each donor-tissue was run separately. SPLASH is considerably more efficient than other approaches, as it utilizes *k*-mers instead of reference alignment and employs closed-form statistics. It is fully parallelized and implemented in a dockerized Nextflow pipeline, making it suitable for massive analyses on high-performance computing clusters. We report the runtime and required memory for each parallelized step to call significant anchors for multiple donor tissues (Suppl. Figure 5), demonstrating that SPLASH calls significant anchors in <40 minutes for a donor-tissue with 400 SS2 cells (provided there is sufficient capacity for full parallelization).

Among called anchors, those classified as splicing had the highest average number of compactors per anchor (Suppl. Figure 6A), consistent with the fact that most genes have only few highly expressed splice variants per tissue (Ezkurdia et al. 2015).

SPLASH+ classified 5.75% (20,891) of anchors as centromeric (19,989,187 reads) (Suppl. Figure 6B; Suppl. Table 2). The human centromere was assembled for the first time in the T2T genome (Altemose et al. 2022); however, as T2T is based on a single cell line, it lacks population-level or single-cell variation. Supporting the limitations of genome alignment, 86% (46,348) of the compactors for centromeric anchors comprising 14% (2,800,038) of their reads failed to align to T2T. Pericentromeric DNA, including human satellite repeat families HSat1-3, is known to be transcribed in certain in vitro and in vivo contexts, but have not yet been studied at single-cell resolution (Altemose et al. 2022). SPLASH+ detected 6,418 anchors containing two consecutive CCATT repeats (or its reverse complement) which defines HSat2. We analyzed the compactors generated for a single anchor containing 2 consecutive CCATT repeats, 166 compactors in donors 1, 2, 4, 7, 8, and 12 (Figure 2A). Compactor sequence diversity for this anchor is extensive as illustrated in the multiway alignment (Figure 2A). 73 compactors (accounting for 53% of total reads assigned to the compactors for this anchor) did not map to T2T by STAR but all compactors did BLAST perfectly to the T2T genome, supporting the idea that the compactors precisely recover true biological sequences. SPLASH+ detected substantial expression variation in multiple cell types including skeletal muscle satellite, mesenchymal, and tongue basal cells that possess proliferative potential (Figure 2A). Basal cells in donor 4 tongue express 23 of the 26 compactors that were found in this donor and tissue (Suppl. Figure 7). The enrichment in proliferative cell populations suggests the hypothesis that expression levels of pericentromeric repeats and replication are linked, as has been explored in limited previous studies (Lu and Gilbert 2007; Probst et al. 2010).

**Figure 2.**
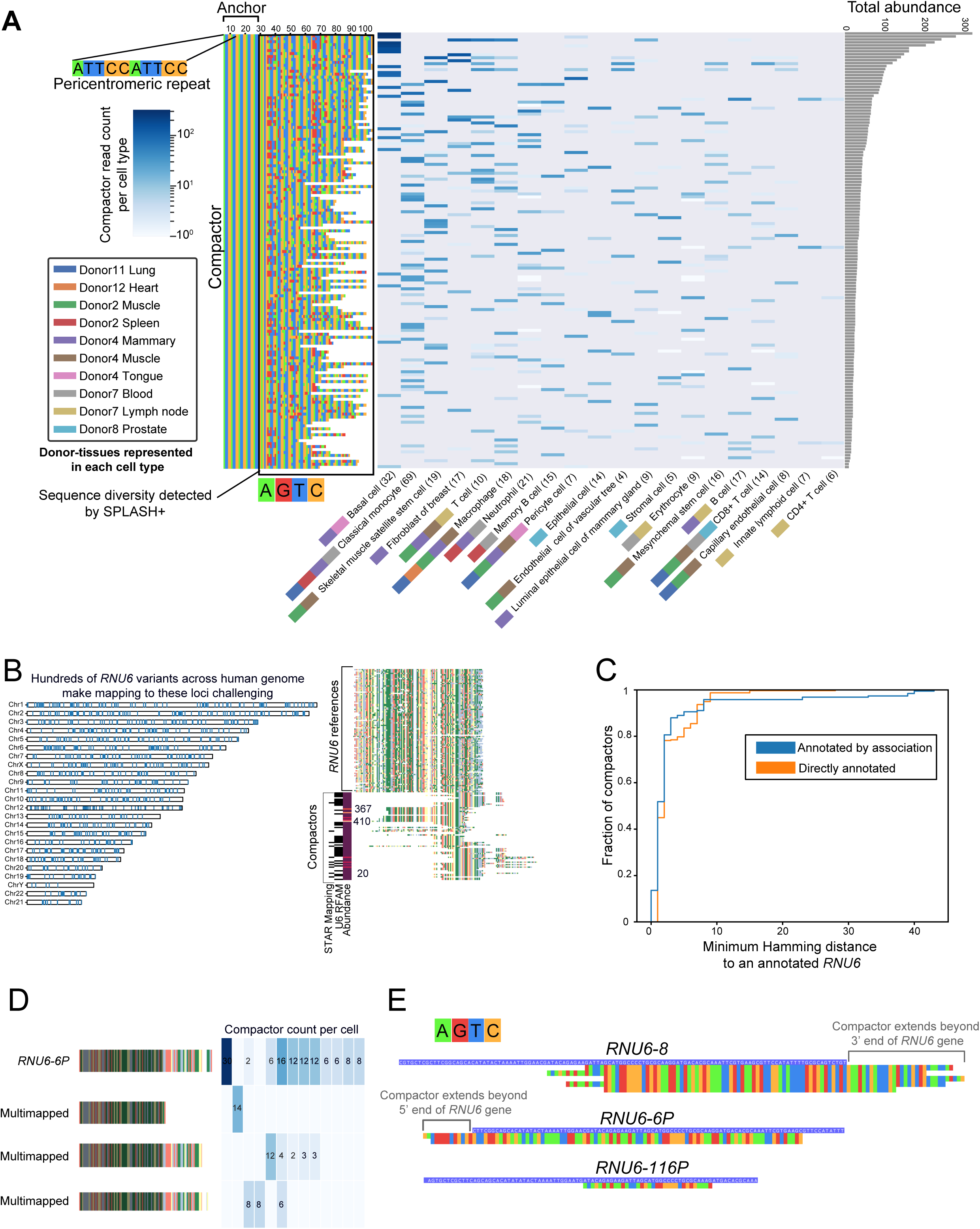
Transcript diversity variation in repetitive RNA loci. (A) SPLASH+ detects variation in centromeric repeat arrays, illustrated for the anchor ATTCCATTCCATTCCATTCCATTCCAC, containing 5 contiguous repeats of ATTCC (the characteristic sequence for canonical pericentromeric). Among the anchors containing ATTCCATTCC, this anchor has the most diverse compactors (147 compactors) ordered through multiway alignment. The heatmap shows the cell-type-specific compactor expression (in logarithmic scale), displaying read counts per cell type (collapsed across donors and tissues). (B) The 71 detected *RNU6* compactors exhibit high conservation with *RNU6* reference sequences, which include 1,281 gene and pseudo-gene loci across the human genome. (C) Direct and annotated-by-association *RNU6* compactors show similar abundances and sequence similarities to *RNU6* reference genes, indicating that the annotated-by-association compactors may be missing from the reference annotation. (D) Heatmap showing differential compactor counts per cell (skin cells from donor 2) for a compactor mapped to the pseudo-gene *RNU6-6P* and other non-uniquely aligned compactors from the same anchor. (E) Multiway alignments of compactors to annotated *RNU6* loci demonstrate alignment both upstream and downstream of annotated boundaries.

We also used SPLASH+ to identify expression variation in non-coding RNA. These loci are extremely difficult to map due to their high copy number and repetitive structure. We bypassed genome alignment and queried compactors against the Rfam (Kalvari et al. 2018), a database of non-coding and structured RNA families, using infernal cmscan (Mistry et al. 2013). The most highly expressed non-coding families were ribosomal RNA in Eukaryotes (6.9M reads, 66% of reads assigned to Rfam-annotated compactors), Prokaryotes (2.1M reads, 20.64%), Archaea (871K reads; 8.28%), and Microsporidia (432K reads; 4.11%) (Suppl. Figure 8, Suppl. Table 3). Some detected rRNA could represent contamination or microbiome composition, as has also been reported by a recent microbial analysis of human single cells (Mahmoudabadi, Tabula Sapiens Consortium, and Quake 2023). The most abundant non-ribosomal noncoding RNA was U6-snRNA (*RNU6*) (22K reads; 0.22% of all Rfam-annotated reads), a small nuclear RNA involved in the spliceosome. *RNU6* has recently been shown to have high cytoplasmic representation (Mabin et al. 2021), suggesting this abundance may be expected due to polyadenylation. More than 75% of compactors assigned to *RNU6* failed to map by STAR, due to multimapping to the 1000+ annotated *RNU6* loci (Figure 2B). We employed the annotation-by-association approach for unaligned compactors associated with anchors with at least one *RNU6*-matching compactor. We computed the minimum Hamming distance for each annotated-by-association compactor and directly-annotated compactor (directly matched an *RNU6* variant by Rfam mapping) relative to the reference set of *RNU6* variants. Both groups had comparable hamming distance to the reference *RNU6* variants (Figure 2C), suggesting that annotated-by-association compactors are potentially false negatives missed by Rfam annotation. There were eight distinct annotated *RNU6* or *RNU6*-pseudogene variants that had uniquely-matching compactors with hamming distance 0, including *RNU6-6P* whose compactors had non-uniform single-cell expression in donor 2 skin (Figure 2D).

We also observed that 30% of *RNU6* compactors had aligned past the 3’ end of the gene, while only 3% mapped upstream of the gene. For example, compactors mapping to *RNU6-8* (659 supporting reads) spanned 45 bps downstream of its 3’ end (Figure 2E). This strongly supports the expression of *RNU6* variants with extended 3’ end, to our knowledge for the first time. There were also *RNU6* variants with compactors mapping upstream of 5’ end (*RNU6-6P, 315* supporting reads) or completely within the annotated locus (*RNU6-116P*) (Figure 2E).

### SPLASH+ improves precision of spliced calls and identifies extensive splicing in *CD47* including novel isoforms

Over 95% of human genes undergo alternative splicing (Pan et al. 2008), but the number of dominant expressed isoforms is mainly based on bulk tissue-level analyses and remains a topic of ongoing debate (Ezkurdia et al. 2015; Arzalluz-Luque and Conesa 2018). This debate arises from alignment-based reference-first approaches with approximate statistical inference due to problems associated with mapping to multiple isoforms (Zheng, Ma, and Kingsford 2022).

Across all donors and tissues, SPLASH+ reported 20,385 anchors (11,995 unique anchor sequences from 3,700 genes) classified as splicing (i.e., anchors with spliced alignment for at least one of their top two compactors), referred to henceforth as splicing anchors, representing single-cell-regulated alternative splicing (Suppl. Table 4). 73.2% of splicing anchors corresponded to annotated alternative splicing junctions (Figure 3A). 1,387 anchors had >10% reads mapping to the unannotated junction of which 706 anchors were found in more than one donor-tissue pair.

**Figure 3.**
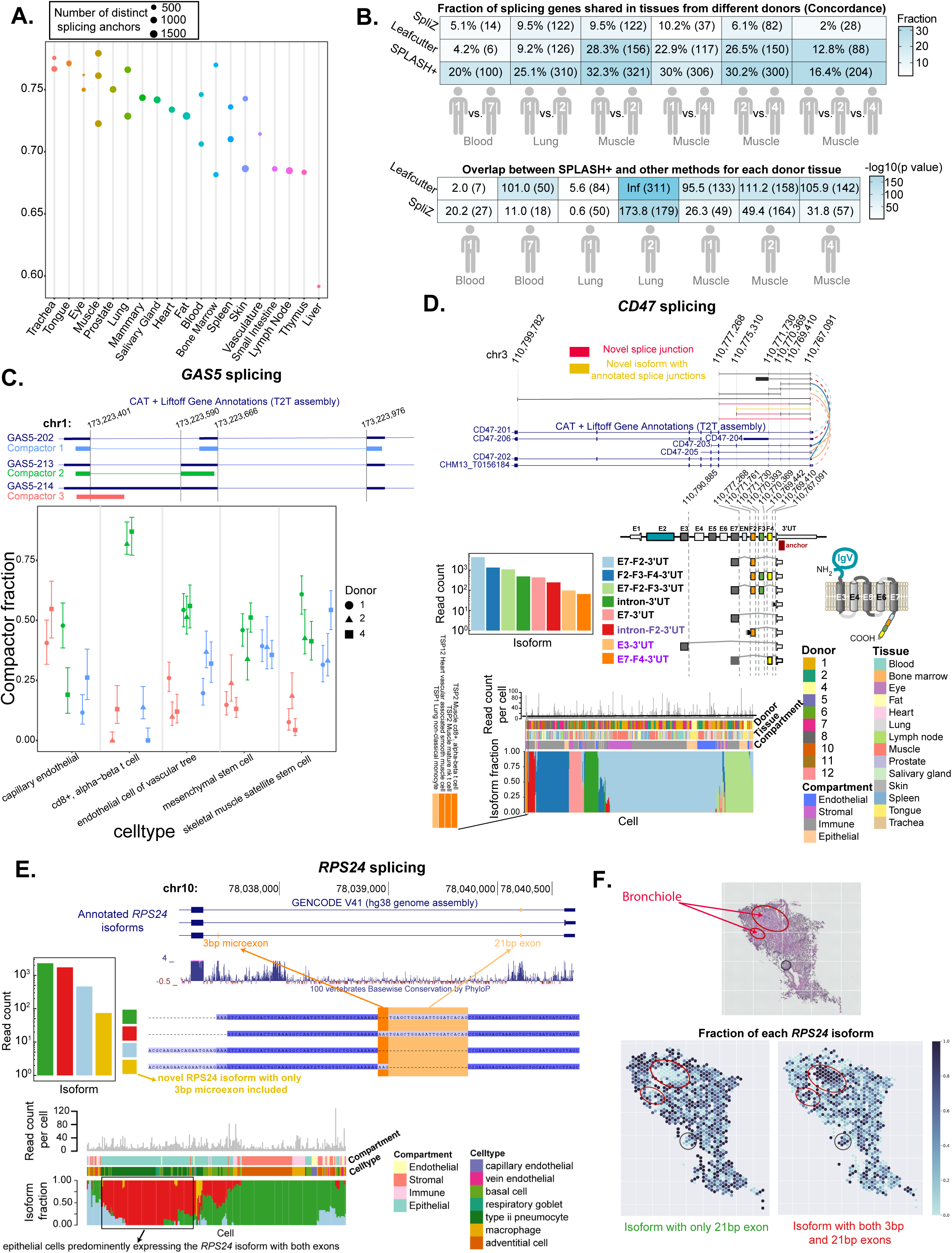
SPLASH+ enables de-novo analysis of alternative splicing in single cells. (A) Dot plot showing the number of distinct splicing anchors and the fraction of them representing annotated alternative splicing per donor-tissue pair, with >55% involving annotated alternative splicing events (based on T2T). Each dot represents a distinct donor corresponding to a specific tissue within the dataset. Muscle has the most number of donors (3 donors). (B) Heatmap (top) showing the concordance of splicing genes (fraction of genes called by a method that are shared in the same tissue from different donors) for SPLASH+, SpliZ (Olivieri, Dehghannasiri, and Salzman 2022), and Leafcutter (Li et al. 2018) in lung, blood, and muscle. This plot suggests that, despite not using cell metadata, SPLASH+ achieves higher concordance in all tissues compared to both Leafcutter and SpliZ. The numbers in parentheses indicate the absolute number of shared genes. Heatmap (bottom) shows the logarithmic binomial p-values (with the numbers in parentheses indicating the absolute number of genes called by both methods) for testing the significance of the number of genes called by both SPLASH+ and Leafcutter, and by both SPLASH+ and SpliZ for each donor tissue. To further emphasize on different cell type composition in tissue replicates, we used different symbols for a tissue from different donors. (C) The reproducible compartment-specific alternative splicing of *GAS5* in muscle cells from 3 donors (intervals are 95% binomial confidence interval), with cd8+, alpha-beta t cells consistently showing a higher inclusion rate for the isoform with shorter intron (green isoform). (D) SPLASH+ detects extensive single-cell expression variation of eight *CD47* isoforms, including novel isoforms, for a single anchor across 10 donors and 14 tissues. The heatmap, sorted by hierarchical clustering, includes a novel *CD47* isoform (yellow) with annotated junctions (isoform coordinates: chr3:110767091:110771730--chr3:110771761:110775311). Cells with >5 reads are included and horizontal bars show the donor, tissue, and compartment identity for each cell. (E) Four alternative splice variants of *RPS24* in lung (from donor 2) involve inclusion/exclusion of ultraconserved cassette exons, including a 3-bp microexon. SPLASH+ detects a novel isoform with only the microexon, missed by both STAR and BLAT, confirmed through multiway alignment. The heatmap shows compartment-specific usage of four *RPS24* isoforms, with the isoform containing the 3-nt and 21-nt exon predominantly expressed in epithelial cells while the novel isoform predominantly expressed in type ii pneumocytes. (F) Spatial validation of *RPS24* alternative splicing using 10x Visium data of lung tissue. Top panels show isoform fractions (left: isoform excluding microexon, right: isoform including microexon). The histology image (bottom) shows bronchial structures (red ellipses), with higher expression of the isoform including the microexon.

We compared SPLASH+ against SpliZ (Olivieri, Dehghannasiri, and Salzman 2022), an algorithm for predicting single-cell-regulated splicing and Leafcutter (Li et al. 2018), a method designed to detect differential splicing in bulk RNA-seq which has also been partially extended to scRNA-Seq. We define splicing genes called by each method in each donor tissue as: genes with cell-type-specific splicing reported by SpliZ (as SpliZ essentially identifies splicing at gene level), genes with at least one splicing anchor reported by SPLASH+, and genes with at least one intron cluster detected by Leafcutter.

We used *Tabula Sapiens (Tabula Sapiens Consortium* et al. 2022)* a scRNA-Seq dataset from primary tissues with no ground truth. Thus, for evaluation, we measured reproducibility of the called splicing across replicates, a standard practice for evaluating discoveries on such datasets. Focusing on the lung (800 cells, 2 donors), blood (538 cells, 2 donors), and muscle (1200 cells, 3 donors) where >1 donor was available and are among the tissues with the most profiled cells, SPLASH+ achieved higher concordance for the same tissue between different donors than both SpliZ and Leafcutter (Figure 3B, top; Suppl. Figure 9 A,B,C). In all 3 tissues, SPLASH+ showed significantly higher reproducibility for splicing genes. For example in blood, SPLASH+ found 20% (100/500) of splicing genes in both donors, whereas SpliZ and Leafcutter found only 5.1% (14/272) and 4.2% (6/143) in both, respectively (Figure 3B). Similarly for lung, SPLASH+ found 25.1% (310/1,235) compared to SpliZ’s 9.5% (122/1,289) and Leafcutter’s 9.2% in both donors (Figure 3B). For muscle, both SPLASH+ and SpliZ called almost the same number of unique genes across three donors. Yet, 204 genes (16.4%) by SPLASH+ were shared among all three donors, while SpliZ and Leafcutter only had 28 and 88 shared genes, respectively (Figure 3B, Suppl. Table 5).

Except for Donor 1 lung and Donor 2 muscle, SPLASH+ called more splicing genes compared to both SpliZ and Leafcutter in all donor tissues (Suppl. Figure 9D). Also, for every donor tissue, the splicing genes called by both SPLASH+ and Leafcutter, and also by both SPLASH+ and SpliZ were more than expected by chance (significant binomial p-values, being less than <10^-6 except for two donor tissues for leafcutter and one donor tissue for SpliZ) (Figure 3B, bottom), providing orthogonal support for the validity of SPLASH+ calls. In summary, SPLASH+ achieved higher concordance than both SpliZ and Leafcutter while identifying more genes as differentially spliced than those methods and showed significant overlap with the calls of those methods. This improvement by SPLASH+ is noteworthy given that it did not use any cell type metadata for inference.

The most highly expressed anchor missed by SpliZ but found by SPLASH+ in all three muscle replicates was noncoding RNA *GAS5*. *GAS5* shows reproducible cell type- and compartment-specific alternative splicing, where cd8+ t cells in all replicates consistently show higher rates for the isoform with shorter intron (Figure 3C) compared to other cell types. Leafcutter detected *GAS5* cell-type-specific splicing for cd8+ t cells only when comparing cd8+ t cells with skeletal muscle satellite stem cells in donor 4, but missed it when comparing cd8+ t cells with four other cell types shown in Figure 3C in donors 1, 2, and 4. We should also note that Leafcutter is blind to intron retention (e.g., the one corresponding to compactor 3) as it only considers split reads. While previous studies have shown opposing effects of *GAS5* splice variants on promoting cell proliferation and apoptosis in tumorigenesis (Mazar et al. 2016; Lin et al. 2022), to our knowledge, this is the first report of the cell-type-specific splicing of *GAS5*.

Recent reference-based metadata-guided studies on human cells and experimental validations have found that *MYL6* and genes within the TPM family (*TPM1*, *TPM2*, *TPM3*) undergo highly cell-type-specific alternative splicing (Olivieri et al. 2021). All these genes were also found by SPLASH+, highlighting its power for detecting cell-type-specific patterns even without utilizing cell type metadata. In muscle, true positives *MYL6* and *TPM1* were significant in all three donors; in contrast, SpliZ only called *TPM1* and *MYL6* in donors 1 and 2. Both SPLASH+ and SpliZ called *TPM2* and *TPM3* in only donor 4.

We also investigated *CD47*, a clinical target for both cardiovascular events (Kojima et al. 2016) and cancer immunotherapy (Gordon et al. 2017) as our previous work showed *CD47* isoform expression was compartment-specific (Olivieri et al. 2021). Among all *CD47* anchors classified as splicing, SPLASH+ detected 10 distinct spliced isoforms, including 5 novel isoforms. One of the *CD47* anchors reveals expression of 8 distinct isoforms including 2 novel isoforms (Figure 3D), all impacting either the cytoplasmic or transmembrane domains. Compartments prefer different isoforms: endothelial and stromal compartments predominantly express E7-F2-3’UT isoform, while immune and epithelial cells in addition to this isoform also express F2-F3-F4-3’UT and E7-F2-F3-3’UT isoforms, respectively. One of the isoforms detected is intron-F2-3’UT (red, Figure 3D); if this isoform indeed represents full intron retention, it would also result in a stop codon after the first transmembrane domain, similar to E7-3’UT isoform. We also detected two novel isoforms: E3-3’UT and E7-F4-3’UT, being the only detected isoforms in 4 cells (Figure 3D). We independently tested for the existence of the novel junctions in these isoforms across all human RNA-seq data deposited in the SRA database (∼285K datasets) via the Pebblescout (Shiryev and Agarwala 2024), a fast k-mer tool that queries RNA-seq studies for the existence of user-provided kmers (Methods). This analysis showed significantly higher prevalence of the two detected novel junctions (E3-3’UT and E7-F4-3’UT) in SRA compared to the control set of unannotated junctions generated by concatenating all exon pairs absent from annotated junctions in *CD47* (Suppl. Figure 10): Pebblescout reported 131K, 9,540, and 2,983 studies on average for the annotated junctions, the two SPLASH-detected unannotated junctions, and the control set, respectively.

SPLASH+ revealed new insights into splicing of *RPS24*, a highly conserved essential component of the ribosome. *RPS24* has 5 annotated isoforms that include ultraconserved intronic sequence and microexons (Olivieri et al. 2021). SPLASH+ detected 4 *RPS24* isoforms in donor 2’s lung cells, consistent with our previous findings of compartment specificity for this gene. Moreover, SPLASH+ identified a novel isoform containing only a 3bp microexon which both STAR and BLAT missed and was detected only through multiple sequence alignment (Figure 3E). Given *RPS24* is highly expressed, we were able to further validate its regulated splicing by analyzing 10x Visium spatial transcriptomic data from lung (SRR14851100). We used *RPS24* anchors detected in scRNA-seq as seeds for compactor generation in the Visium data. We found significant spatial organization for the two compactors corresponding to the two most abundant *RPS2*4 isoforms in scRNA-seq. The isoform containing the microexon had higher expression in the two bronchiolar structures marked by red ellipsis in the histology image (Figure 3F), comprising mostly club and ciliated cell types (Travaglini et al. 2020). Both cell types exhibited concordant enrichment of microexon containing isoform in both lung replicates in scRNA-seq, confirming our spatial predictions (Suppl. Figure 11). Together, this analysis shows that without using any cell metadata or reference, SPLASH+ can generate biologically consistent predictions for splicing in human tissue.

### Genes with pan-tissue, single-cell regulated splicing are enriched for splicing factors and histone regulation

2,118 genes with splicing anchors called by SPLASH+ in more than one tissue, including 10 genes found in 18/19 tissues, being referred to as *core genes* (Figure 4A). We performed GO enrichment analysis on 61 genes with splicing anchors detected in at least 15 tissues. Remarkably, enriched pathways with the highest log-fold change were all involved in mRNA processing and splicing regulation (Fisher test, FDR p-value < 0.05, Figure 4B). These results imply that splicing factors and histone modifications themselves are under tight splicing regulatory mechanisms in diverse tissues, possibly co-regulating their expression.

**Figure 4.**
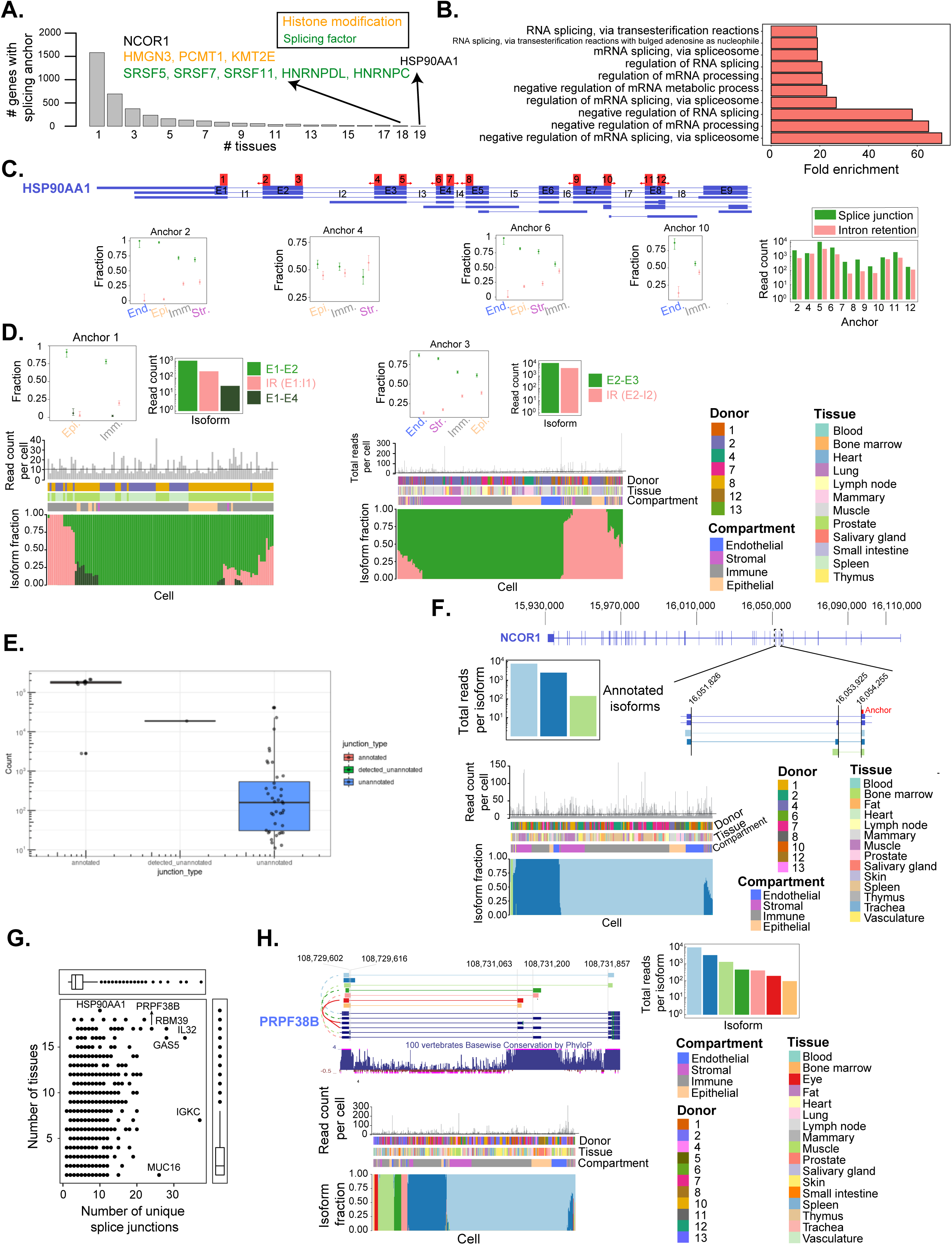
Pan-tissue alternative splicing regulation of splicing factors and histone modifications. (A) 57% of genes (2,118) with significant splicing in at least 2 tissues, including 10 genes found in >= 18 tissues. Notably, 8 splicing factors (green) and histone modifications (brown) are present. (B) GO enrichment analysis of genes found in >15 tissues reveals pathways related to mRNA processing and splicing regulation (Fisher test, FDR corrected p-value< 0.05). (C, D) Pan-tissue intron retention diversity in *HSP90AA1*, the only SPLASH+ core gene found in all 19 tissues. Twelve anchors detect differential intron retention events in 7 (out of 8) *HSP90AA1* introns. The compartment-specific expression of compactors corresponding to intron retention (pink) and splice junctions (green); the barplot on the right shows total read counts for intron retention and splicing isoform for each anchor. Fractional isoform distribution is depicted for the anchor with the most compactors (anchor 1) and the anchor found in the most tissues (anchor 3). (E) Number of SRA studies containing each *HSP90AA1* junction reported by Pebblescout supports SPLASH+’s novel splice prediction. Junctions are grouped as annotated, unannotated, and one detected unannotated junction (between exons E1 and E4). (F) Single-cell dependent alternative splicing of gene *NCOR1,* involving 3 different splicing isoforms for the *NCOR1* anchor found in the most tissues. (G) Plot showing the number of tissues and unique splice junctions for each gene with a splicing anchor. *IL32*, *RBM39*, and *IGKC* have the most unique splice junctions. (H) Alternative splicing of *PRPF38B* involving an intron retention and six splicing isoforms, showing one of the most diverse isoform structures with significant alternative splicing in 11 donors and 17 tissues.

SPLASH+ reveals diverse splicing regulation in the 10 core genes, detecting 71 isoform variants, including four unannotated isoforms, one in each of the *HNRNPC*, *KMT2E*, *SRSF7*, and *SRSF11* genes. While each of the core genes are known to have significant regulatory roles, the extent and complexity of their splicing regulation, as revealed by SPLASH+, has been underappreciated.

*HSP90AA1* was the sole core gene found in all 19 tissues (Figure 4C). *HSP90AA1* is one of two isoforms of the HSP90 heat shock protein functioning in myriad cellular processes including a chaperone of protein folding (Hoter, El-Sabban, and Naim 2018) and transcriptionally regulated under cell stress (Zuehlke et al. 2015). We detected anchors with 12 distinct differential intron retention events for 7 introns of this gene including unannotated intron retention events for introns 1 through 4 and intron 7, and a novel splicing between the first and fourth exons (Figures 4C,D). Detected intron retentions are highly compartment-specific, with higher expression fractions for immune and stromal cells compared to epithelial and endothelial cells (Figures 4C,D). For 9 anchors, immune cells had the highest intron retention fraction compared to other cells. Anchor 6 had the strongest differential pattern between compartments with 44%, 22%, 17% and 0% for intron retention in immune, epithelial, stromal, and endothelial cells, respectively. Due to its known transcriptional regulation upon stress, we cannot exclude the possibility that the regulated intron retention is due to differential compartment-specific response to dissociation stress. However, compartment-specificity and abundance of intron retention forms suggest *HSP90AA1* has previously-unknown post-transcriptional regulation, even if part of the detected signal is differentially regulated physiological response to dissociation. We also queried the novel junction (between exons E1 and E3) via Pebblescout, which indicated a 6-fold increase in prevalence compared to the control set of unannotated junctions (Figure 4E). To our knowledge, splicing regulation in *HSP90AA1* represents a potentially novel mechanism to tune the protein levels of this critical molecular chaperone.

Five core genes (50%) are themselves splicing factors, including nuclear ribonucleoproteins (hnRNPs) *HNRNPC*, *HNRNPDL* (Suppl. Figure 12), and SR family members *SRSF5*, *SRSF7*, *SRSF11*. The detected compactors represent complex isoforms, some un-annotated, and some including splicing into ultraconserved intronic regions known to create poison exons in *SRSF5*, *SRSF7*, *SRSF11*, and *HNRNPDL* (Lareau et al. 2007; Königs et al. 2020; Raihan et al. 2019; Ni et al. 2007). The remaining four core genes are involved in histone regulation or nuclear co-repression: *KMT2E* a histone methyltransferase with known mutations in neurodevelopmental disorders (O’Donnell-Luria et al. 2019), *PCMT1*, another histone methyltransferase (Biterge et al. 2014), *HMGN3*, a high mobility group nucleosome-binding protein and transcriptional repressor and *NCOR1*, the nuclear co-repressor (Perissi et al. 2010). SPLASH+ detects a poison exon and intron retention event in *NCOR1* (Figure 4F) and *KMT2E,* respectively, which are predicted to trigger nonsense mediated decay. Core genes all have portions of highly conserved intronic sequences, suggesting a mechanism for splicing regulation. Together, these results support the idea that alternative isoforms play a critical regulatory role that includes use of premature stop codons and complex alternative splicing in the 5’ UTR, gene body, and 3’ UTRs. While these isoforms had been predicted by analysis of EST data and in cell culture, to our knowledge, direct evidence of regulated splicing patterns of any of these regulators in single cells has been previously missing (Ding et al. 2022; Lareau et al. 2007).

We also used SPLASH+ to identify genes classified as splicing with high numbers of variants, which are known to drive organization of complex tissues (Schmucker et al. 2000; Yagi 2008). We identified genes with the most unique splice junctions across the entire dataset (Figure 4G): *IL32*, *GAS5*, and *RBM39* (33, 28, and 28 junctions, respectively). Additionally, 49 genes, including *PRPF38B*, *TACC1*, *CCDC66*, and *TAX1BP3*, had at least 15 distinct splice junctions (Figure 4G). Among these genes are known oncogenes *TACC1* (Cully et al. 2005) and *CDC37* (Gray et al. 2008) with 19 and 11 splice variants, respectively. Splicing factors *SRSF10* and *RBM39* (each found in 17 tissues) were also highly ranked, having 16 and 28 splice variants, respectively, and are all associated with tumor initiation and growth (Kim et al. 2015; Xu et al. 2021; Shkreta et al. 2021). For *PRPF38B*, a splicing factor with prognostic biomarker potential in breast cancer (Abdel-Fatah et al. 2017), 17 distinct splice junctions were detected (across all of its anchors) in 17 tissues. One of its anchors shows compactors with complex alternative splicing involving skipping of two cassette exons, alternative 5’ splice sites, intron retention, and a novel splice junction, which is the dominant isoform in 4 immune and stromal cells (Figure 4H). SPLASH+ reveals complexity of splicing regulated at the single cell level missed by current methods, and supports the idea that many human genes have single-cell-specific splicing patterns, rather than exclusively favoring a dominant form.

### SPLASH+ rediscovers and expands the scope of V(D)J transcript diversity

Single cells can somatically acquire copy number variation, SNPs, or repeat expansions. Detection of genetic diversity in single cells has required custom experimental and computational workflows. Our statistical analysis demonstrates that under the null assumption where all cells in a donor express only two alleles of any given splice variant, the probability of observing many counts for compactors beyond the two genuine ones decays rapidly (Methods). Positive controls expected to violate the null include mitochondria where genomes are polyploid (Barrett et al. 1983) or the rearrangement of immunoglobulin loci in B cells and T cell receptor loci which undergo V(D)J recombination.

To investigate global trends, for each anchor, we collected all distinct compactors across donor-tissues (Suppl. Table 6). Anchors mapping to immunoglobulin genes had the highest number of distinct compactors and differentiated from other genes on this purely numerical criterion (Figure 5A). For example, an anchor mapping to immunoglobulin kappa chain (*IGKC*) had the highest number of compactors (140) across all donors and tissues. As expected, these anchors were observed only in immune tissues. Centromeric anchors defined as containing CCATTCCATT repeats also had substantial compactor diversity (Figure 5A), consistent with the fact that centromeres have substantial sequence diversity within and across individuals.

**Figure 5.**
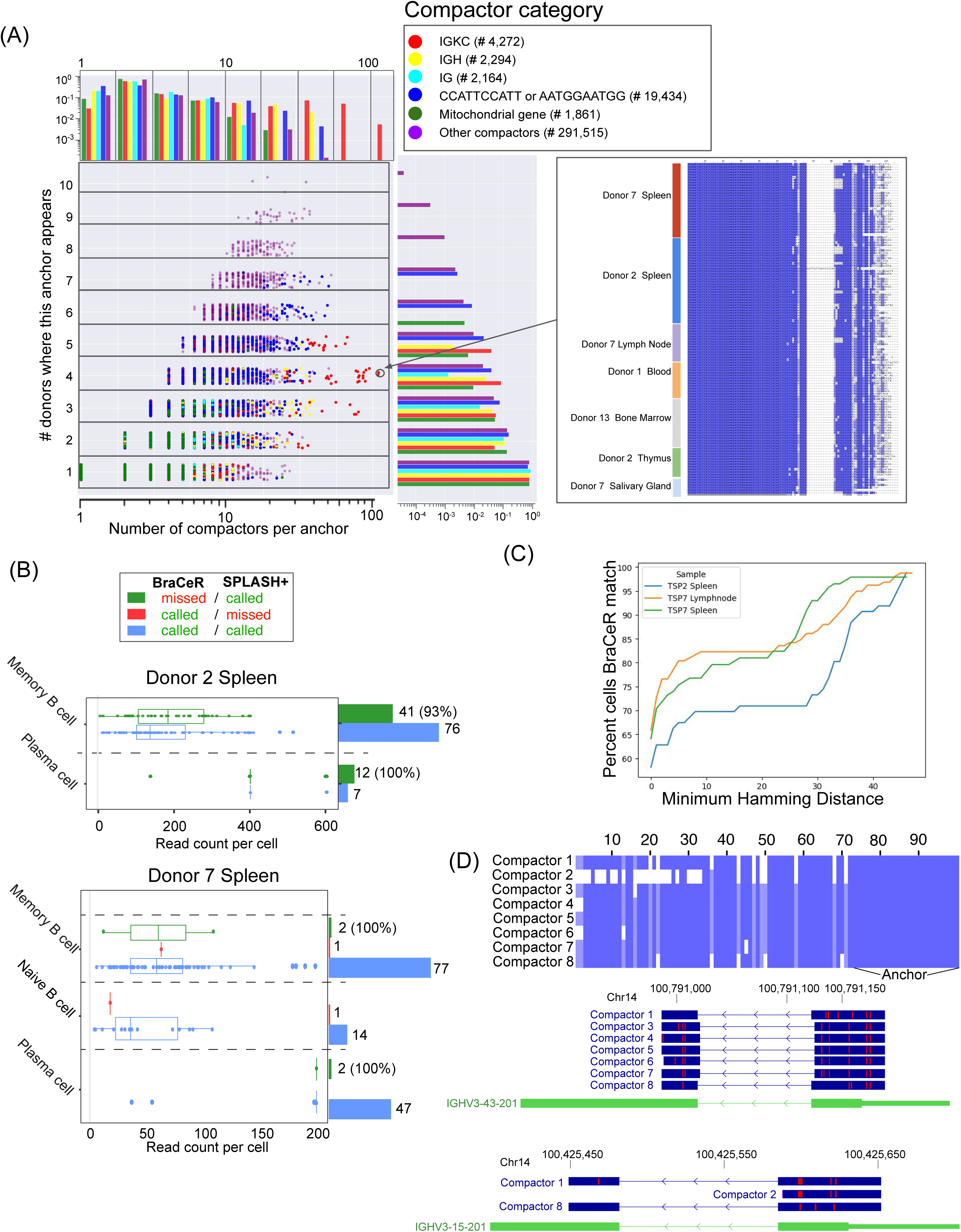
SPLASH+ detects V(D)J recombination with highest levels of transcriptome diversity and achieves higher sensitivity than BraCeR. (A) Plot showing the number of compactors and donors for each anchor, indicating that anchors with the most compactor diversity extend known biology of V(D)J recombination. The top marginal histogram shows the probability that each compactor category falls into a specific range of number of compactors per anchor. The right marginal histogram shows the probability of each category as a function of the number of donors. Multiple sequence alignment of the anchor mapping to an immunoglobulin gene with the most distinct compactors. (B) Comparison of SPLASH+ and BraCeR for spleen cells (that are analyzed by both methods) from donors 2 and 7 shows higher sensitivity for SPLASH+. Cells are called by BraCeR or SPLASH+ based on the presence of at least one BraCeR contig or IG-compactor, respectively. The x-axis shows the highest count for an IG-compactor in each cell and we also show the percentage of cells called only by SPLASH+ (green) within each cell type having at least one in-frame V(D)J transcript as confirmed by IgBLAST. (C) Cells called by both SPLASH+ and BraCeR increase with the minimum Hamming distance between IG-compactors and BraCeR contigs per cell. (D) Alignment of multiple compactors from the same anchor to *IGHV3* shows split mapping to *IGHV3-43-201* and *IGHV3-15-201*.

Mitochondrial genes also had substantial diversity, notably in an anchor from gene *MT-ND5* in donor 1 lung that had 24 compactors. This was the highest number of compactors for a single anchor mapping to an annotated gene within a single donor-tissue. *MT-ND5* is a component of the transmembrane electron transport chain in mitochondria with previously reported recurrent mutations with clinical significance (Jaberi et al. 2020; Wang et al. 2022).

While current approaches require mapping to the genome on a read-by-read basis, SPLASH+ enables a statistics-first micro-assembly to detect variants in the B cell receptor (BCR) locus avoiding genome alignment. We compared SPLASH+ power to detect BCR rearrangement in B cells from spleen of donors 2 and 7 against the state-of-the-art custom pipeline BraCeR (Lindeman et al. 2018). We name a compactor as an “IG-compactor” if STAR maps it to an immunoglobulin gene (Methods). As a more stringent criterion, we further annotated IG-compactors using IgBLAST (Ye et al. 2013) and considered only those IG-compactors that had both variable (V) and joining (J) immunoglobulin gene segments identified through IgBLAST (Methods). In donor 2 spleen, not only did SPLASH+ detect an IG-compactor with annotated V and J genes in every cell for which BraCeR reported a BCR contig, but it also found BCR evidence in 53 additional B cells (41 memory B cell and 12 plasma cell) which BraCeR missed (Figure 5B; Suppl. Table 7). Similarly, in donor 7 spleen both BraCeR and SPLASH+ found evidence of BCR rearrangement in the same 47 plasma B cells, but SPLASH+ also found BCR evidence in 2 additional plasma B cells (Figure 5B). SPLASH+ missed one cell among memory B cells and naive B cells from donor 7, which BraCeR called. We further investigated SPLASH+ calls missed by BraCeR to see if they had evidence of fully functional in-frame transcripts. 100% of plasma cells in both donors and 93% and 100% of memory B cells in donors 2 and 7, respectively, that were called only by SPLASH+ had at least one in-frame IG-compactor as predicted by IgBLAST (Figure 5B), suggesting that higher sensitivity of SPLASH+ can result in the identification of high-confidence fully-functional immunoglobulin transcripts with implications in adaptive immune response.

Of the cells analyzed by both algorithms, SPLASH+ called IG-compactors in 136 cells from donor 2 spleen, and 142 and 123 in donor 7 spleen and lymph node (Suppl. Table 7). We tested if SPLASH+’s IG-compactors were concordant with BraCeR’s calls in these cells by computing the minimum Hamming distance between the two sets of BraceR contigs and SPLASH+ IG-compactors for each cell (Figure 5C). A high fraction of cells had perfect matches between SPLASH+’s and BraCeR’s sequences: 58.1%, 65.8%, and 64.1% for donor 2 spleen, donor 7 spleen, and donor 7 lymph node, respectively, with increasing concordance as a more relaxed minimum Hamming distance criterion is considered between IG-compactors and BraceR contigs (Figure 5C). To identify IG-compactors that were missed by BraCeR, compactors with Hamming distance of greater than 30 to all BraCeR contigs were collected. 416 anchors had more than 3 of such compactors. One of these anchors had 8 compactors in donor 2 spleen where each compactor had partial alignment to one or both of two different IGHV loci, likely representing distinct V segment inclusion (Figure 5D). In summary, SPLASH+ automatically detects V(D)J rearrangement, agreeing with and extending what is detected by BraCeR in expected B cell subtypes, with implications for downstream biological inference and opportunities to explore other sequences nominated by SPLASH+ that do not meet the stringent criteria used here.

### Cell type-specific hypermutation or RNA editing including in intronic regions of *AGO2*, UTRs of *ANAPC16* and the 5’ and translational start of *ARPC2*

We investigated anchors that had either comparable or more distinct compactors across donor-tissues than those associated with V(D)J. This list includes anchors with compactors showing abundant canonical Adenosine-to-inosine (A-to-I) RNA editing in *ANAPC16* (Figure 6A), a regulator of anaphase, and *AGO2* (Figure 6B), the argonaut protein involved in miRNA targeting.

**Figure 6.**
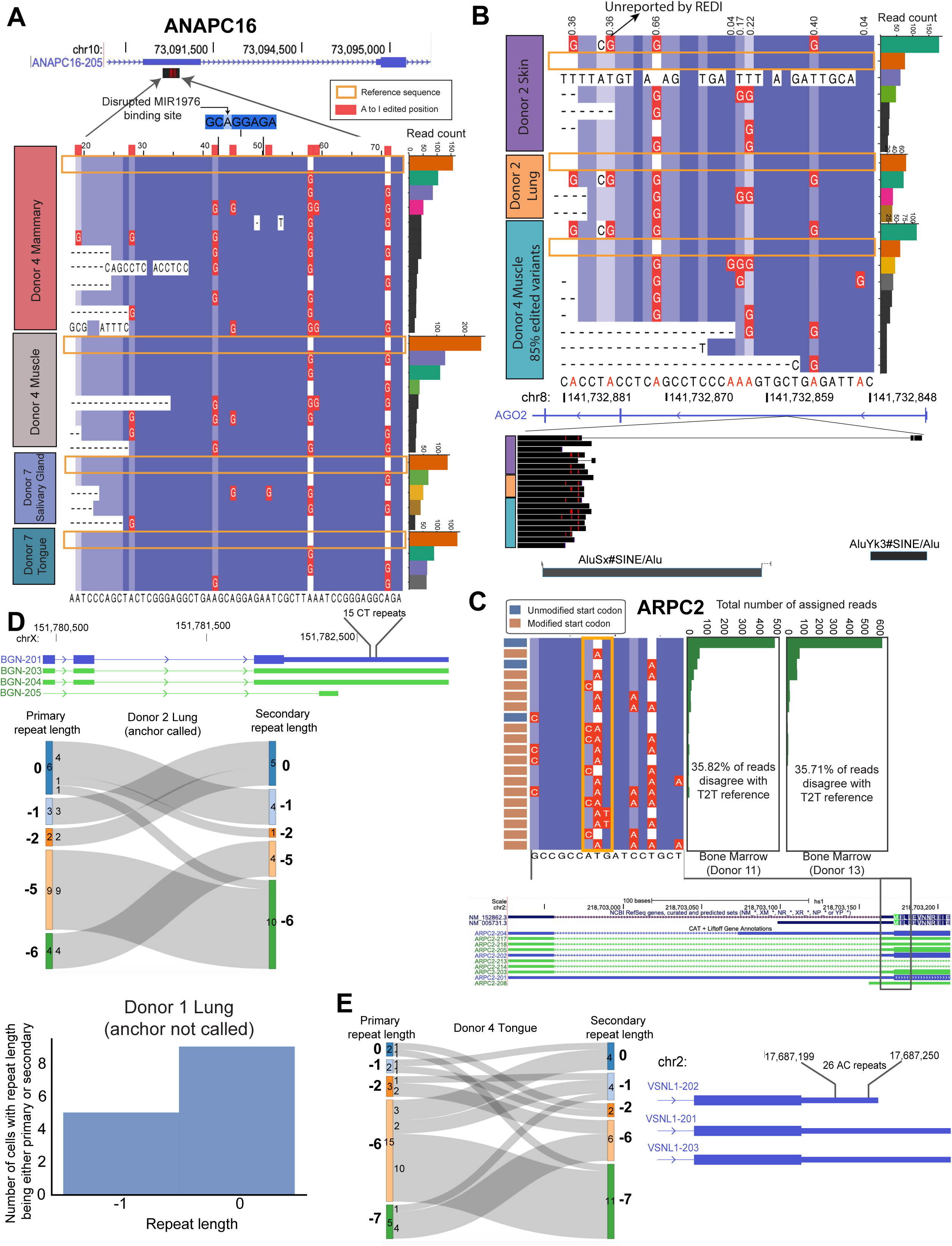
Single-cell-regulated RNA editing and repeat polymorphism detected *de novo* by SPLASH+. (A) Multiple sequence alignment of compactors generated *de novo* by SPLASH+ reveals extensive cell-type regulated editing in 5’ UTR of *ANAPC16*. Red indicates A-to-I edits, and compactors matching the reference allele for each donor-tissue are highlighted in orange boxes. Side bar plots show total counts for each compactor in each donor tissue, with top 4 compactors in each donor tissue color-coded by shared sequences across donor tissues. Predicted miRNA binding site (Chen and Wang 2020) in *ANAPC16* is disrupted by observed edits in all four donor-tissues. (B) Similar analysis for *AGO2* identifies RNA editing within an Alu element including a circular RNA in the third compactor from donor 2 skin. (C) *ARPC2* sequence diversity shows 20 distinct variants in a 17-bp region (positions 58-74 in the compactor sequences). Multiple sequence alignment reveals base-pair changes between the 5’ UTR and translational start (chr2: 218,703,167–218,703,183; T2T assembly). All kmer counts from donor bone marrow exceed expected sequencing error rate by orders of magnitude. (D) Sankey plot displays cell counts for primary (repeat with the highest count in cell) and secondary (repeat with the second highest count in cell) repeat lengths in donor 2 lung (donor tissue in which the anchor was called). The reference repeat length is indicated as 0 and repeat contractions are indicated relative to this reference length. Bar plot shows cell counts with reference or one repeat contraction. Each cell is counted for a repeat length if either its primary or secondary repeat matches that length. (E) Similar analysis for repeat polymorphism in *VSNL1* shows cell counts for each combination of secondary and primary repeat lengths.

In *AGO2*, a critical enzyme for RNA interference, SPLASH+ generated 18 compactors from a single anchor across skin and lung from donor 2 and muscle from donor 4 (Figure 6B). Similar to *ANAPC16*, the majority of reads were assigned to edited variants, constituting 84%, 64%, and 85% of reads in skin, lung, and muscle, respectively. These compactors support A-to-I canonical intronic editing in an Alu element at 8 positions, including one editing, supported by 36% of reads across the three donor-tissues, that is not reported in the REDIPortal database of RNA editing events in humans (Picardi et al. 2017). Also, a compactor representing 15% of reads in donor 2 skin was both edited and showed circular RNA backsplice junction (Figure 6B), suggesting that splicing precedes editing. The extent of editing in these loci and extremely low support from a comprehensive, ultra-deep reference database (each editing event being supported by at most 14 studies out of 9,642 studies curated in REDI) suggest reads at this locus would be unmapped or mismapped with conventional pipelines (Eisenberg and Levanon 2018). In fact, as shown in the BLAT alignment of these compactors (Figure 6B), for the first and fifth compactors in donor 2 skin, BLAT missed a few editing events and instead reported a split alignment, illustrating the challenges of reference-based alignment method for detecting these events.

Extensive RNA editing diversity was also found in *ARPC2*, the actin-related protein 2/3 complex subunit 2. We focused our analysis on the single anchor from this gene with the most compactors in a single donor (16 compactors). Compactors for this anchor represent prevalent base pair changes with respect to the reference, with consistent editing rates (∼36%) in bone marrow cells from donors 11 and 13 (Figure 6C). These editing events lack a known mechanistic explanation as they are neither canonical A-to-I editing nor show signatures of reverse-transcriptase induced base pair changes due to RNA base modifications (Werner et al. 2020). Intriguingly, the changes are concentrated in the start codon and would likely affect translation initiation. *ARPC2* has other known non-canonical translation regulation: an internal ribosome entry site in its 5’ UTR (Al-Zeer et al. 2019), and un-annotated splicing in its 5’ UTR, suggesting the possibility of non-canonical translation initiation. Because of its surprising nature, we tested if apparent editing in *ARPC2* existed in other donors, but SPLASH+ was underpowered to detect it. We generated compactors using the above anchor for all cells in the study: 15% of all reads in the dataset containing this anchor had discrepant bases in the 17-bp window (chr2: 218,703,167–218,703,183), and 11% had base pair changes in the start codon, much higher than expected by chance under sequencing error rate (median error rate .01% for the Illumina NovaSeq 6000) (Stoler and Nekrutenko 2021).

We further used Pebblescout to test the frequency of editing events detected in *ARPC2* (Methods). Editing events detected by SPLASH+ were reproducibly found in human RNA-seq samples (Suppl. Figure 13A). To investigate if the detection rate is higher than base pair changes due to sequencing error, we also constructed and queried decoy sequences containing base pair changes not predicted by SPLASH+. The detection rate was significantly higher for the editing locations predicted by SPLASH+ (Suppl. Figure 13B) and the prevalence difference between editing and decoy sequences grew with the number of base pair changes. For example, k-mers with 2 detected edits by SPLASH+ were present in an average of 26K samples whereas decoys were detected in only 163 (∼160-fold change). No decoys with 3 edits were detected whereas SPLASH+ kmers were detected on average in ∼3K samples.

The reproducibility across donors, tissue specificity, stereotyped positions, and the level of diversity are strong evidence against these base-pair changes being an artifact or arising in DNA. In summary, SPLASH+’s automatic statistical inference identifies extensive and novel editing in single cells. To our knowledge, these events have not been and cannot be detected with current custom workflows (Cohen-Fultheim and Levanon 2021).

### Evidence for repeat polymorphism including in *BGN* and *VSNL1*

Other anchors with high compactor diversity show evidence of repeat polymorphism. For example, for an anchor mapping to 3’ UTR of *BGN*, compactors show multiple AG dinucleotide repeat lengths (Figure 6D). *BGN* codes for biglycan and has roles in metabolic pathways and cell proliferation (Ying et al. 2018; Morimoto et al. 2021). Dinucleotide repeats are known to be polymorphic, but repeat length variation could also be generated during PCR, through a process called slippage (Shinde 2003). Thus, we investigated if polymerase slippage could explain the repeat polymorphisms found by SPLASH+. The profile of *in vitro* Taq PCR slippage has been studied, showing that after each PCR cycle the repeat element is contracted with a non-negligible probability *p* (Shinde 2003). Under this error model, the contraction process is equivalent to a binomial process and therefore we should observe a unimodal probability distribution of repeat lengths shorter than the reference repeat in the genome.

The top two dominant repeats in the vast majority of cells in donor 2 lung were of different lengths and distinct from the reference repeat (Figure 6D; Methods): 9 cells and 10 cells had contractions of 5 and 6 as their primary and secondary repeat, respectively. We tested if observed repeat variants in *BGN* are consistent with the slippage error model. In donor 2 lung, the dominant repeat lengths were 15 (reference repeat length) and 10 (20% and 53% of reads, respectively), inconsistent with being generated *in vitro* by PCR slippage, which would generate a unimodal probability distribution of repeats ranging from 15 to 10. Instead, this supports a model that donor 2 has two *BGN* alleles with 15 and 10 repeats. To further test if these variable repeat lengths could be explained by PCR artifacts, we generated compactors in donor 1 lung where SPLASH+ did not call the *BGN* anchor. The expression patterns for two repeat lengths in Donor 1 were compatible with being generated due to PCR artifacts as the reference allele comprised majority of reads (78%), while contractions of 1 or 2 AG dinucleotides accounted only for 22% of reads which is in contrast to what was observed in donor 2 lung. This, together with the donor-specific repeat polymorphism is strong evidence that SPLASH+ calls *BGN* because single cells express different allelic repeat lengths, perhaps due to allelic imbalance, rather than being due to PCR artifacts.

Another repeat polymorphism found by SPLASH+ was in *VSNL1* (Visinin like 1 protein), a neuronal sensor calcium protein (Figure 6E). Prior literature shows that repeat polymorphism in *VSNL1*, is highly conserved in vertebrates and implicated in dendritic targeting (Ola et al. 2012; Riley and Krieger 2009). Contractions of 6 and 7 repeats were most abundant (together 75% of reads, Figure 6E); Contractions of 0, 1 and 2 repeats had 8%, 7%, and 10% of reads, respectively, with no intermediate repeats between these and contractions of 6 and 7. Highlighting the importance of avoiding cell type metadata for testing, *VSNL1* polymorphism was detected predominantly in one cell type: tongue basal cells, which are thought to be stem cell progenitors (Iwai et al. 2008). If these alleles were due to polymerase slippage, error models predict observing other contractions such as −8; however, none were observed. This, along with the diverse single-cell expression of non-allelic repeat variants, suggests donor-specific somatic diversification of repeat numbers, likely originating from somatic variation within a donor rather than PCR.

## Conclusion

Alignment to the human reference genome is often considered a prerequisite for the analysis of RNA-sequencing, and great efforts have been made to provide complete and curated reference genomes and transcript annotations. Similarly, cell type metadata is currently viewed as a critical starting or ending point for the analysis of single-cell sequencing experiments. In this work, we demonstrated that novel aspects of transcriptome regulation can be discovered using a direct statistical approach to analyze sequencing data without cell type metadata, while using reference genomes only for post-inferential interpretation.

In addition to computational and conceptual unification, SPLASH+’s reference-free approach predicts biology in single cells that has been missed by customized bioinformatics methods in multiple domains. SPLASH+ enables automatic discovery of cell-specific diversity in non-coding RNAs, such as *RNU6* and centromeric repeats, which, to our knowledge, are not captured with current bioinformatic approaches. In domains where custom algorithms exist, such as detection of RNA splicing, V(D)J recombination, or RNA editing, SPLASH+ unifies and extends discovery. SPLASH+’s findings of complex splicing regulation in splicing factors provide direct evidence of and extend previous predictions of such regulation from EST databases and DNA sequence. We also uncovered novel cell-regulated splicing in diverse genes, including noncoding RNAs such as *GAS5*, suggesting new candidates to prioritize for functional studies. The extent of splicing diversity uncovered by SPLASH+ further supports the idea that transcriptome complexity in primary single cells is extensive. This implies that a data science-driven approach will be needed to predict regulation and function for this transcriptome diversity, as experimental approaches cannot be scaled to the throughput required to study each isoform.

This manuscript also demonstrates how SPLASH+’s approach unifies disparate areas beyond splicing discovery, including variation in noncoding RNA loci, centromeres, detecting genomic insertions such as Alus, V(D)J recombination, RNA editing, and repeat polymorphisms. This suggests further avenues for discovery of human disease biology in both RNA-seq and DNA-seq where SPLASH+ allows repeat polymorphisms to be further scrutinized. For example, dinucleotide repeats detected in this study are predicted to be bound by CUG-binding protein (*MBNL1*) and TDP-43 (Takahashi et al. 2000; Buratti and Baralle 2001). The repeat polymorphisms identified by SPLASH+ further suggest the potential for predicting cell-specific impacts of repeat expansions, including their contribution to stress granule formation and disease (Sproviero et al. 2017; Estany et al. 2007). To focus this work, we did not include a discussion of other dimensions of transcript diversity found by SPLASH+, including alternative polyadenylation within human transcripts, cell-level variation in indels, potential structural rearrangements within the human genome, and even non-human sequences found by SPLASH+ in this dataset, which include an enrichment of bacteriophages that may reflect the prokaryotic contribution to the human metatranscriptome. We should note that while SPLASH+ is applicable to diverse genomics problems, it may not be suitable for studies where the focus is on differential gene expression analysis. Also, providing better interpretation for those SPLASH+’s calls that do not have BLAST hits is part of the ongoing work.

In this first unbiased systematic analysis of human transcriptomic diversity in single cells, SPLASH+ establishes a unified statistics-first approach to sequence analysis, which reveals prevalent transcript diversity variations overlooked by current bioinformatics tools. While examples discussed in this study provide a glimpse into its complexity due to subsampling of human cells and tissues, analysis of larger single-cell datasets, as well as DNA sequencing, hold potential for uncovering a new generation of genetic and transcriptomic diversity mechanisms as predictors of cellular phenotype or disease. Indeed, SPLASH+ is versatile and applicable to any RNA-seq or DNA-seq study, offering opportunities for a large-scale, statistically driven study of transcriptomes.

## Acknowledgments

We thank all members of the Salzman lab for comments, Aaron Straight, Pragya Sidhwani and Charles Limouse for reading and interpretation of the results on centromeric repeats. Lu Chen for discussion of RNU-6 variation, Liana Lareau for comments. We also thank Serana Tan for her help with interpreting Visium data. We would like to thank Stanford University and the Stanford Research Computing Center for providing computational resources and support that contributed to these research results. J.S. is supported by the Stanford University Discovery Innovation Award, National Institute of General Medical Sciences grants R35 GM139517 and the National Science Foundation Faculty Early Career Development Program Award no. MCB1552196. RB was funded by the NCI grant 5F31CA243170-02. TZB was funded in part by the Stanford Graduate Fellowship and the NSF GRFP.

## Competing Interests

K.C., T.Z.B., and J.S. are inventors on provisional patents related to the original implementation of SPLASH. The authors declare no other competing interests.

## Figures

**Suppl. Figure 1.**
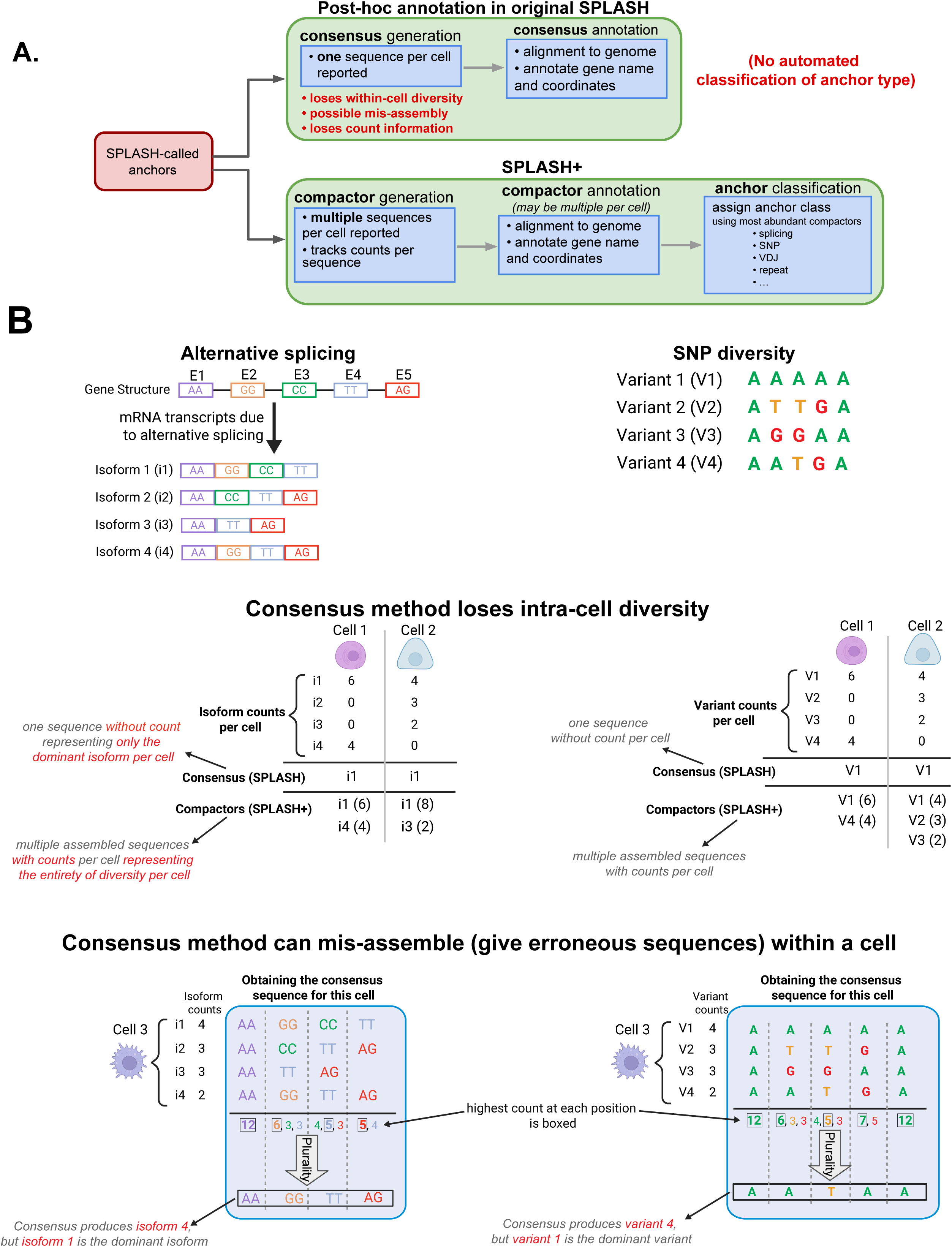
(A) In SPLASH, a single consensus sequence is constructed for each sample (cell) by taking the plurality of bases at each position across targets. This approach loses both within-cell diversity and count information, as it reports only one sequence and may even lead to misassembly. The consensus sequence is then aligned to the genome for gene name and alignment information. Notably, SPLASH lacks automated anchor classification, limiting its application for targeted studies on specific events such as alternative splicing. In contrast, SPLASH+ employs a branching approach to construct long assembled sequences (compactors) with assigned counts for each sample. The most abundant compactors for each anchor are then used to assign biologically relevant events (e.g., alternative splicing, V(D)J recombination) to each anchor. (B) Consensus (used in original SPLASH) fails to capture sequence diversity variation across cells as it produces only one sequence per cell. For example, for the given gene with 4 transcript isoforms (each exon is assumed to have two nucleotides as shown), the consensus sequence for both Cells 1 and 2 is isoform 1 (i1), the dominant isoform in both cells, implying no diversity variation between these cells. However, Cells 1 and 2 have different count distributions of isoforms 2, 3, and 4. For Cell 3, the consensus approach incorrectly represents isoform 4, which has the lowest count. Similarly when sequence diversity is due to SNPs, the consensus approach might fail to show diversity variation as seen with Cells 1 and 2, where both are represented by variant V1 despite their different distributions for other variants. In Cell 3, the consensus incorrectly represents a non-existent variant. We show the details of how the consensus sequence is produced for Cell 3 (for both splicing and SNP diversity).

**Suppl. Figure 2.**
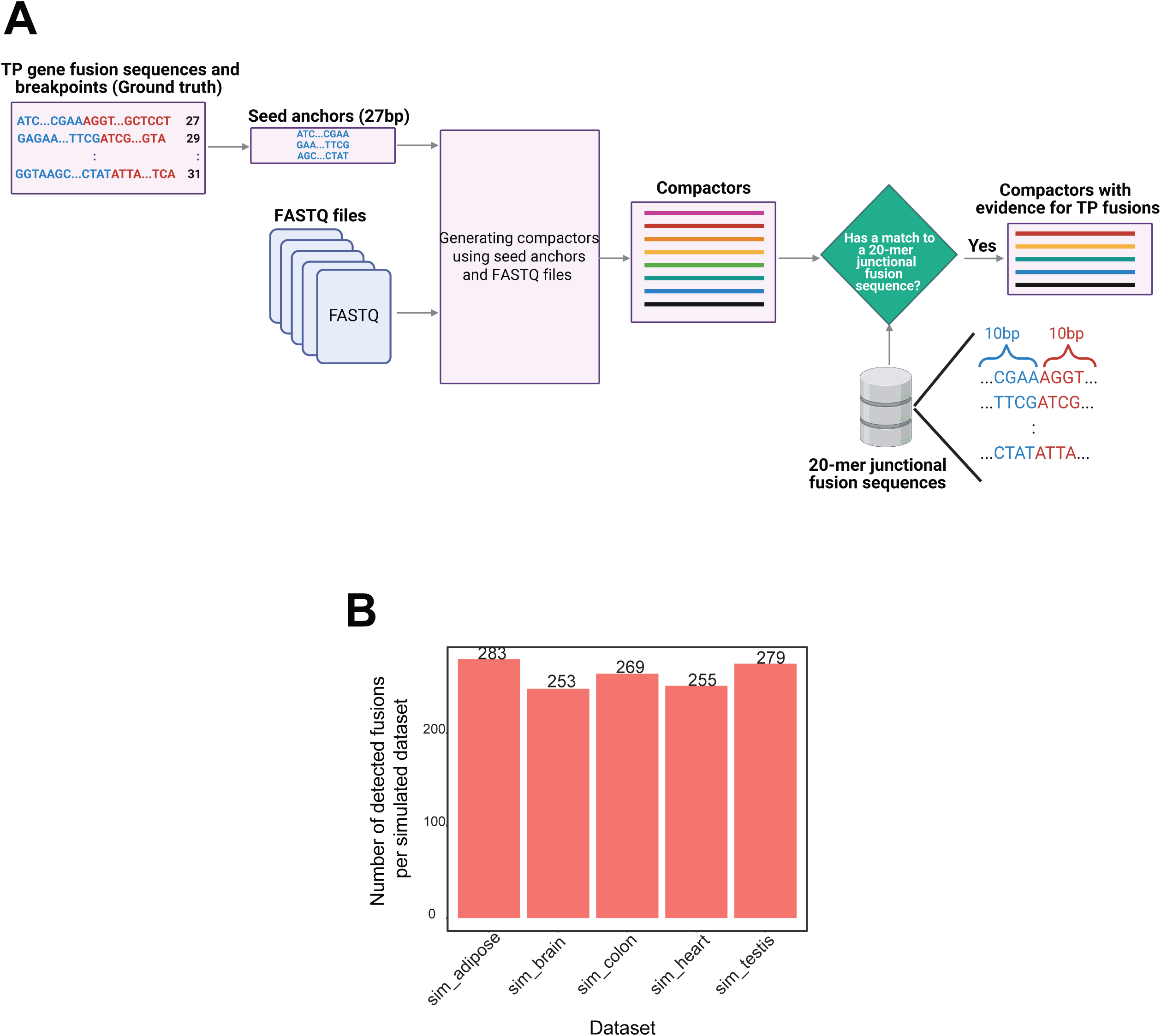
(A) Overview of compactor benchmarking analysis using the fusion benchmarking dataset. The dataset provides the sequence and breakpoint position for each true positive gene fusion. We constructed seed anchors from the 27 base pairs immediately upstream of each breakpoint and generated compactors using these seed anchors using FASTQ files. Each compactor was assigned to a true positive gene fusion if it matched the 20-mer junctional sequence for that fusion, which was created by taking 10 base pairs on each side of the gene fusion junction. For each dataset, the number of unique gene fusions with associated compactors is counted, representing the number of detected fusions reconstructed by compactors for that dataset. (B) Number of detected fusions per simulated datasets. In total 57.8% (1339/2315) of fusions were detected by compactors.

**Suppl. Figure 3.**
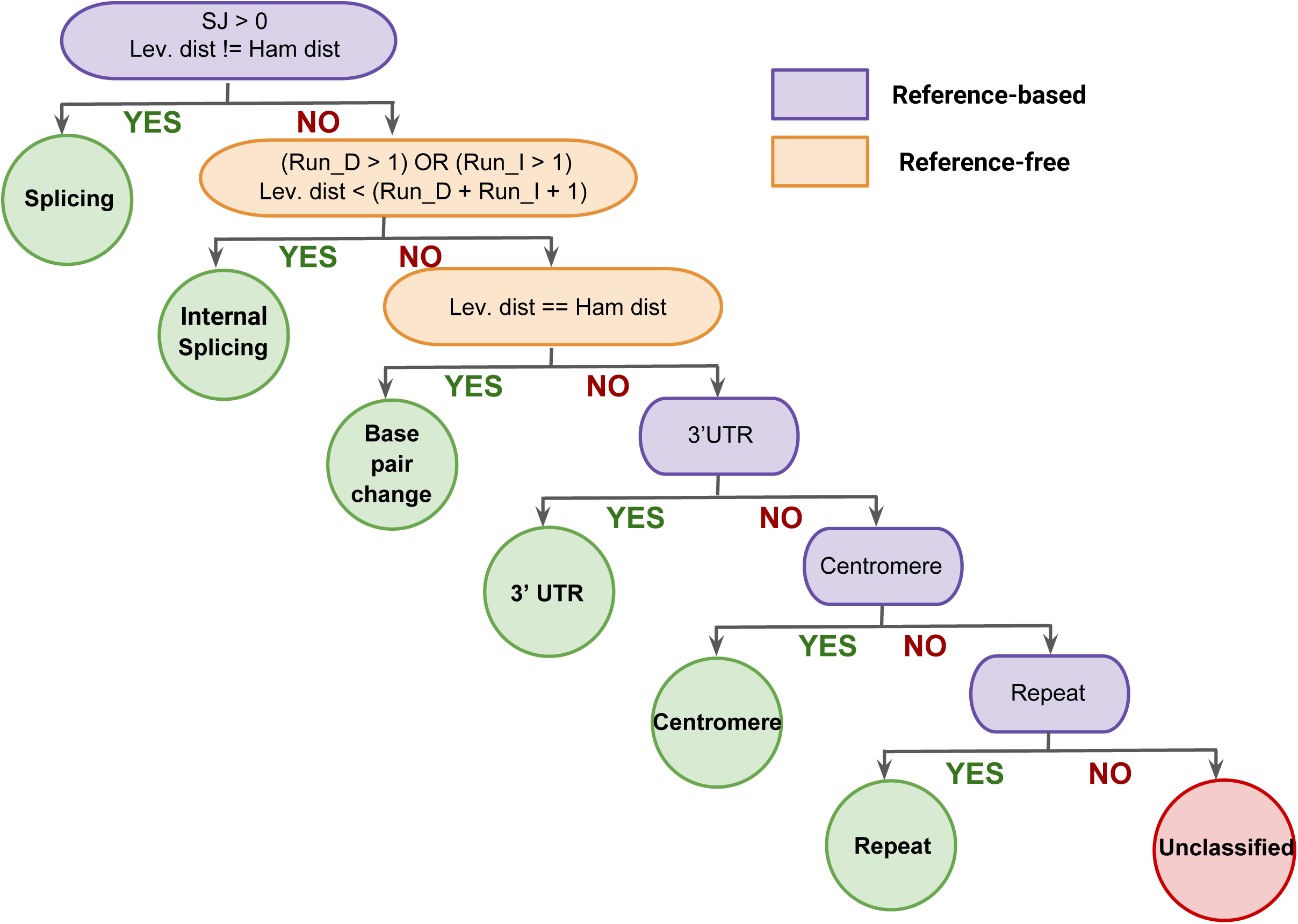
Post-facto classification assigns distinct biologically relevant RNA events to each anchor based on its compactors’ characteristics. Six categories are considered: splicing, internal splicing, base pair change, 3’ UTR, centromere, and repeat (ordered by priority). An anchor that cannot be assigned to any of these categories is referred to as unclassified. Reference-free categories include internal splicing and base pair change; and splicing, 3’ UTR, centromere, and repeats are assigned based on compactors’ alignment to the genome.

**Suppl. Figure 4.**
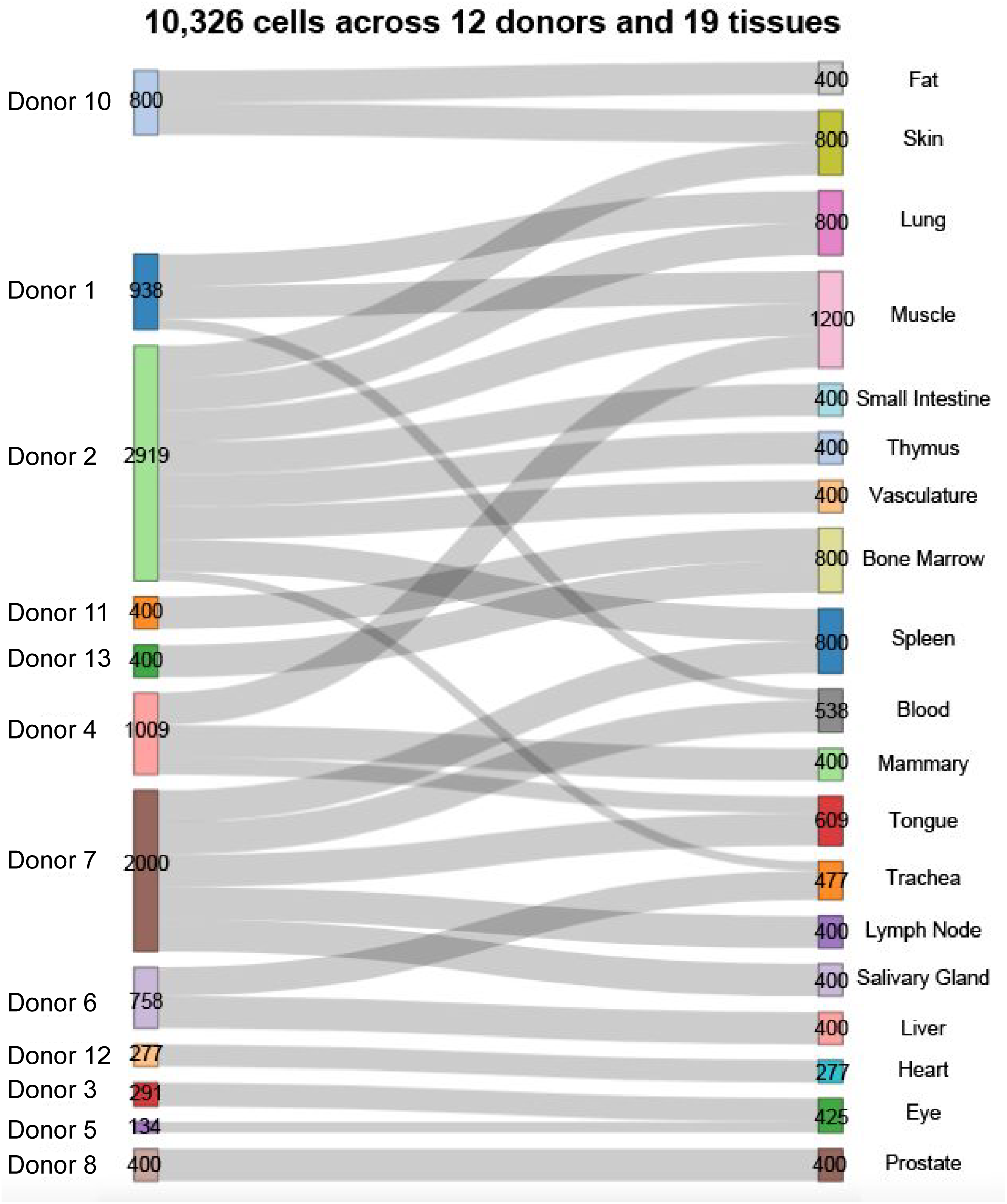
SPLASH+ was run on 10,326 SmartSeq2 cells from 12 donors and 19 tissues (29 donor-tissue pairs), including replicates for several tissues from multiple donors, enabling reproducibility analysis.

**Suppl. Figure 5.**
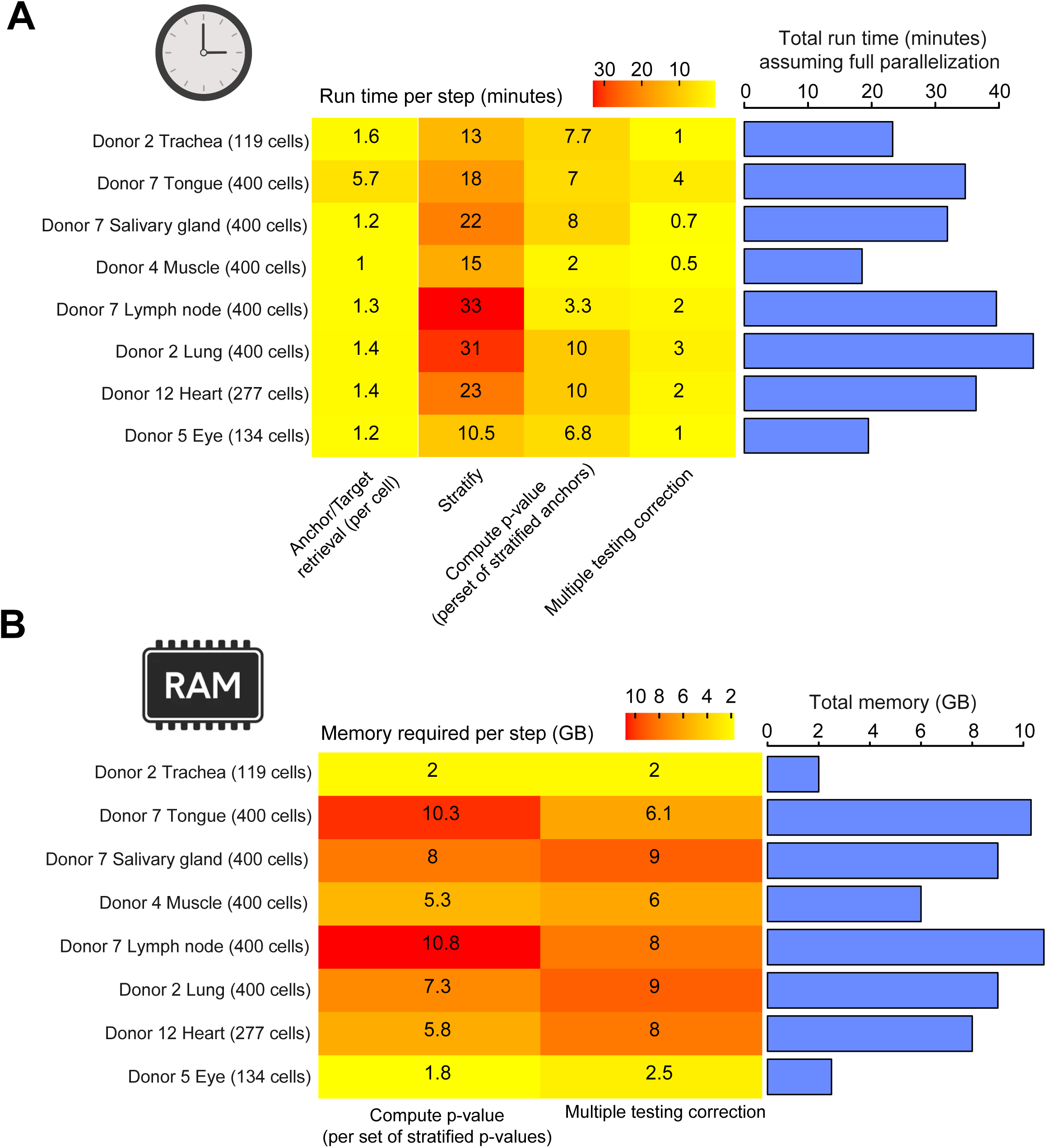
(A) Run time and (B) memory usage per step (extracting and counting anchor and target sequences from input sample in parallel for each input sample, stratification of anchor sequences to allow for parallelization of p-value calculation, p-value calculation in parallel for each set of stratified anchors, and multiple testing correction) in SPLASH across eight donor tissues. SPLASH, being fully parallelized, allows the initial step of parsing input reads and extracting anchor and target sequences to run concurrently for each cell. Thus, the run time for this step is presented for individual cells. Memory requirements are displayed for the two steps consuming the most memory.

**Suppl. Figure 6.**
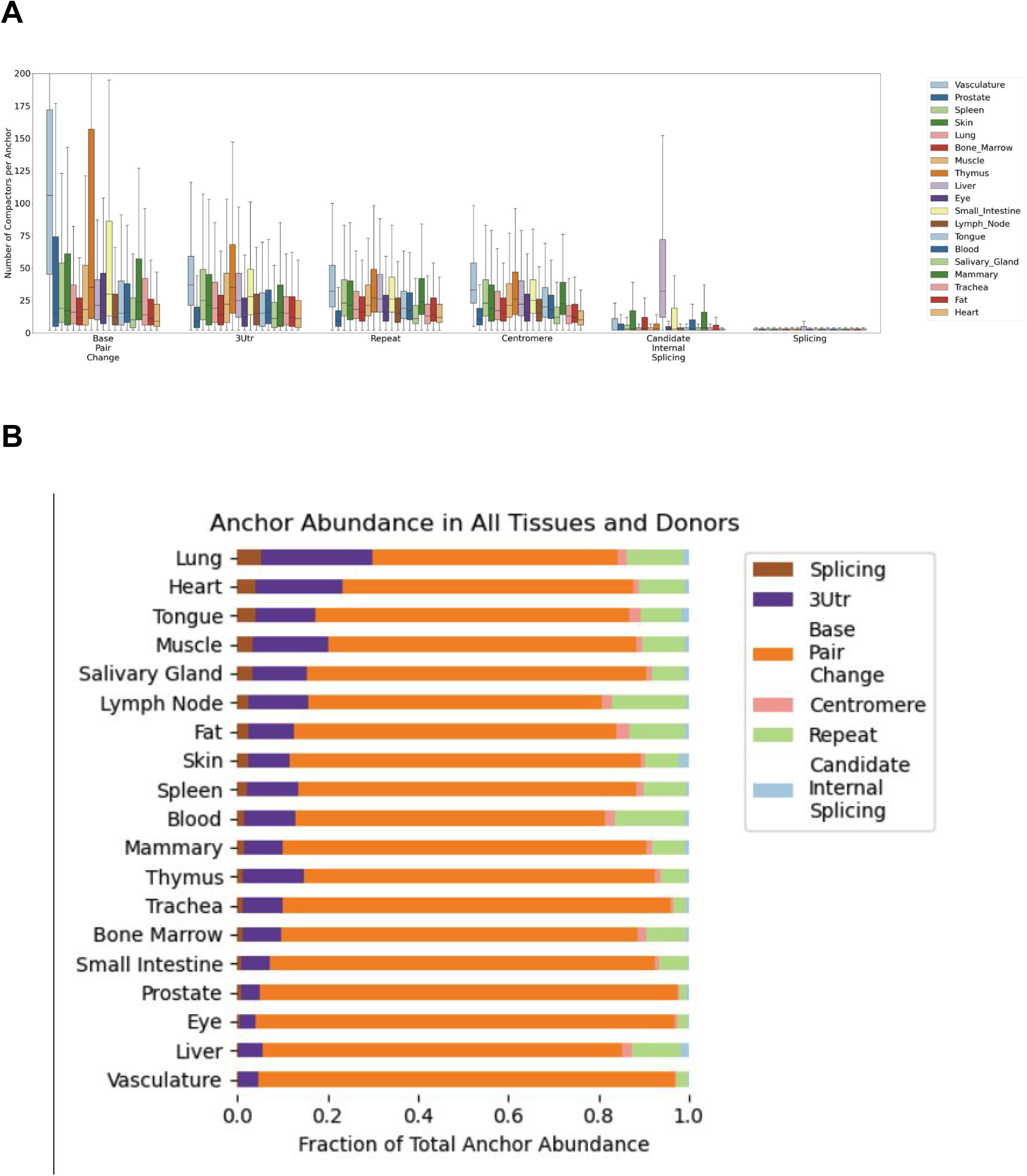
(A) Compactor count per anchor in each tissue, categorized by anchor class, with splicing anchors exhibiting the lowest compactor diversity. (B) Fraction of reads assigned to each anchor category in each tissue, with base pair change anchors having the highest fraction of reads.

**Suppl. Figure 7.**
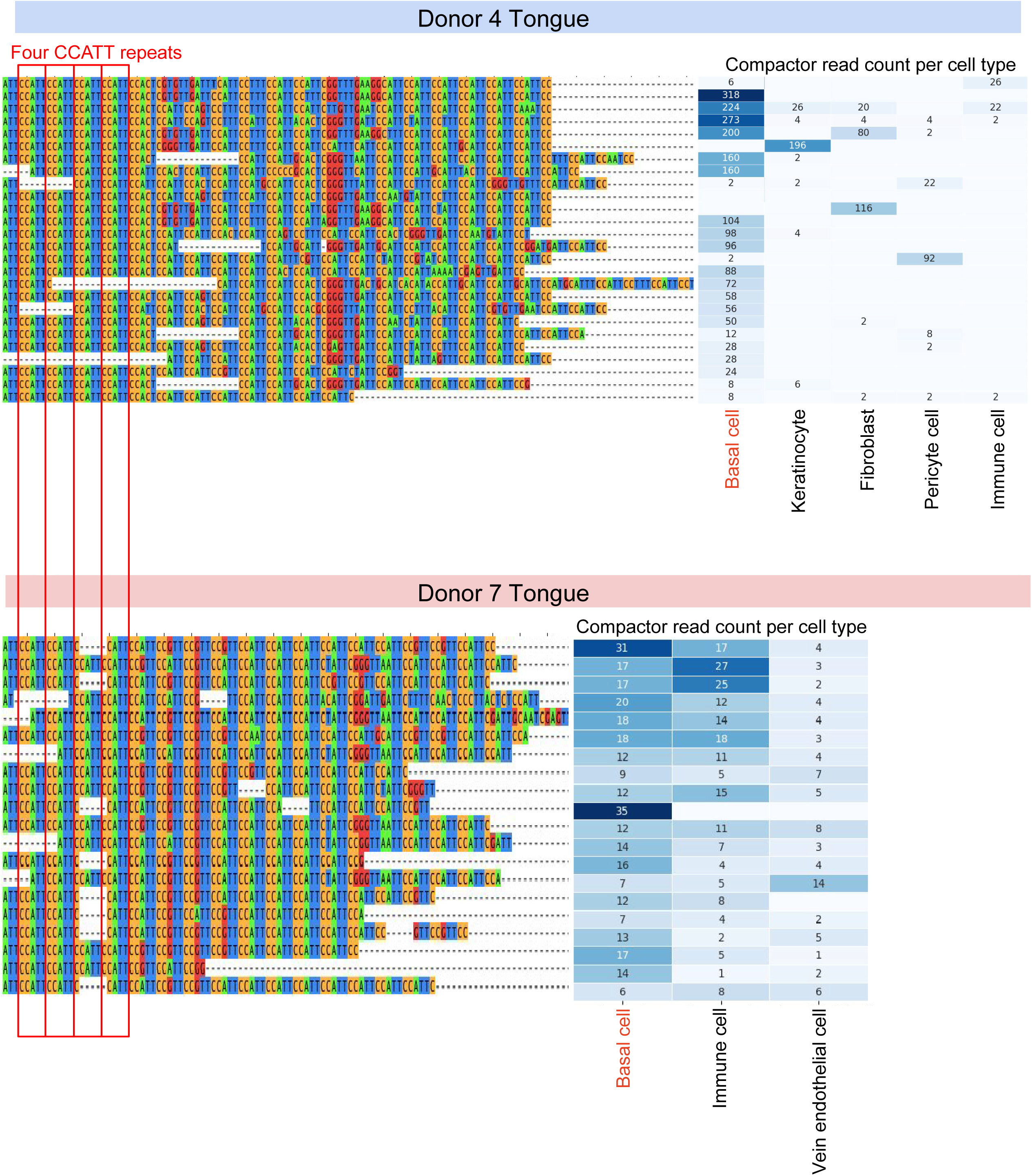
Single-cell-regulated expression of the anchor containing pericentromeric repeat CCATT in tongue replicates (donors 4 and 7). Tongue basal cells consistently exhibit the highest expression in both replicates.

**Suppl. Figure 8.**
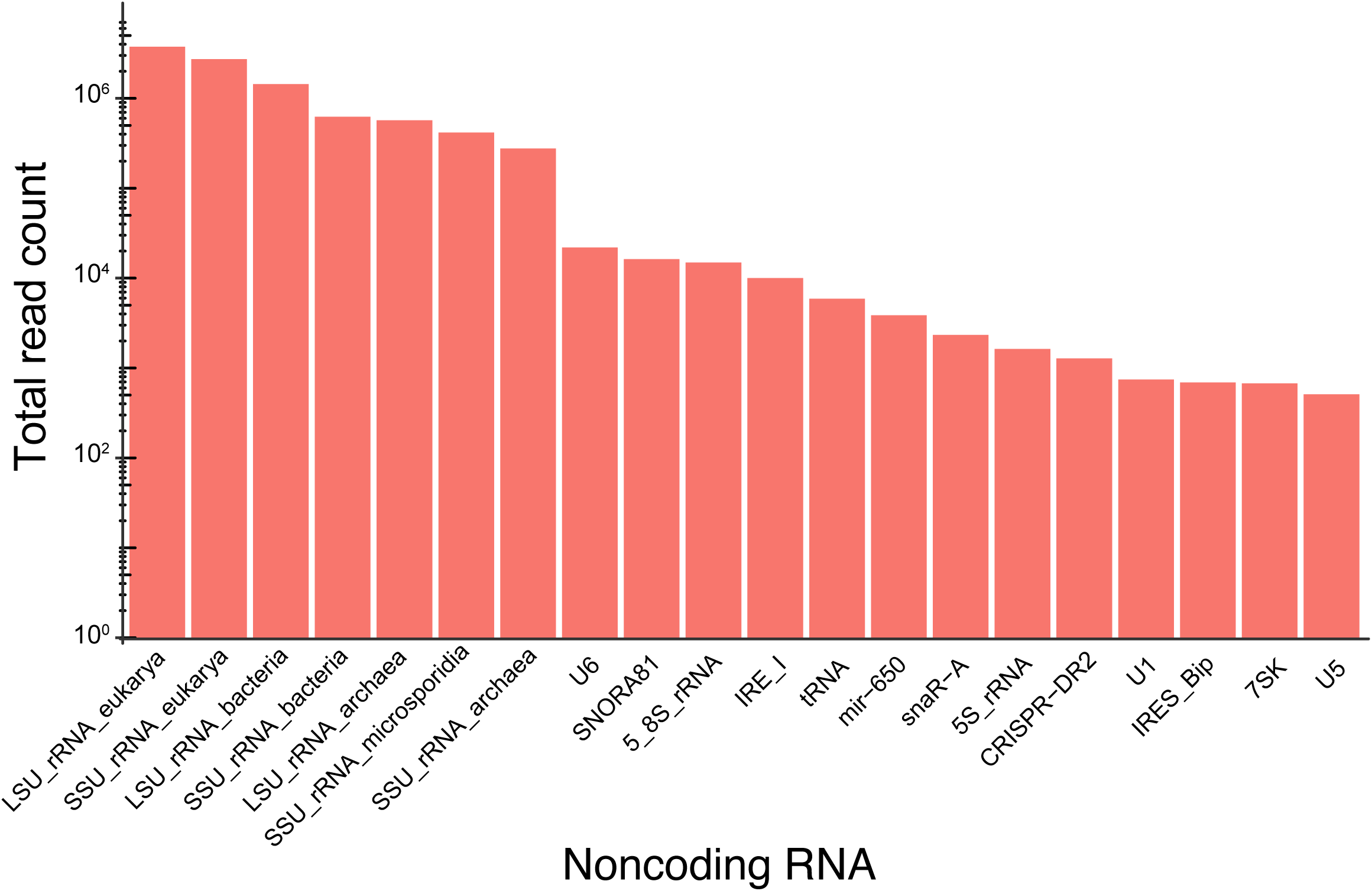
The barplot showing the total read count across the entire dataset for each category of noncoding RNAs.

**Suppl. Figure 9.**
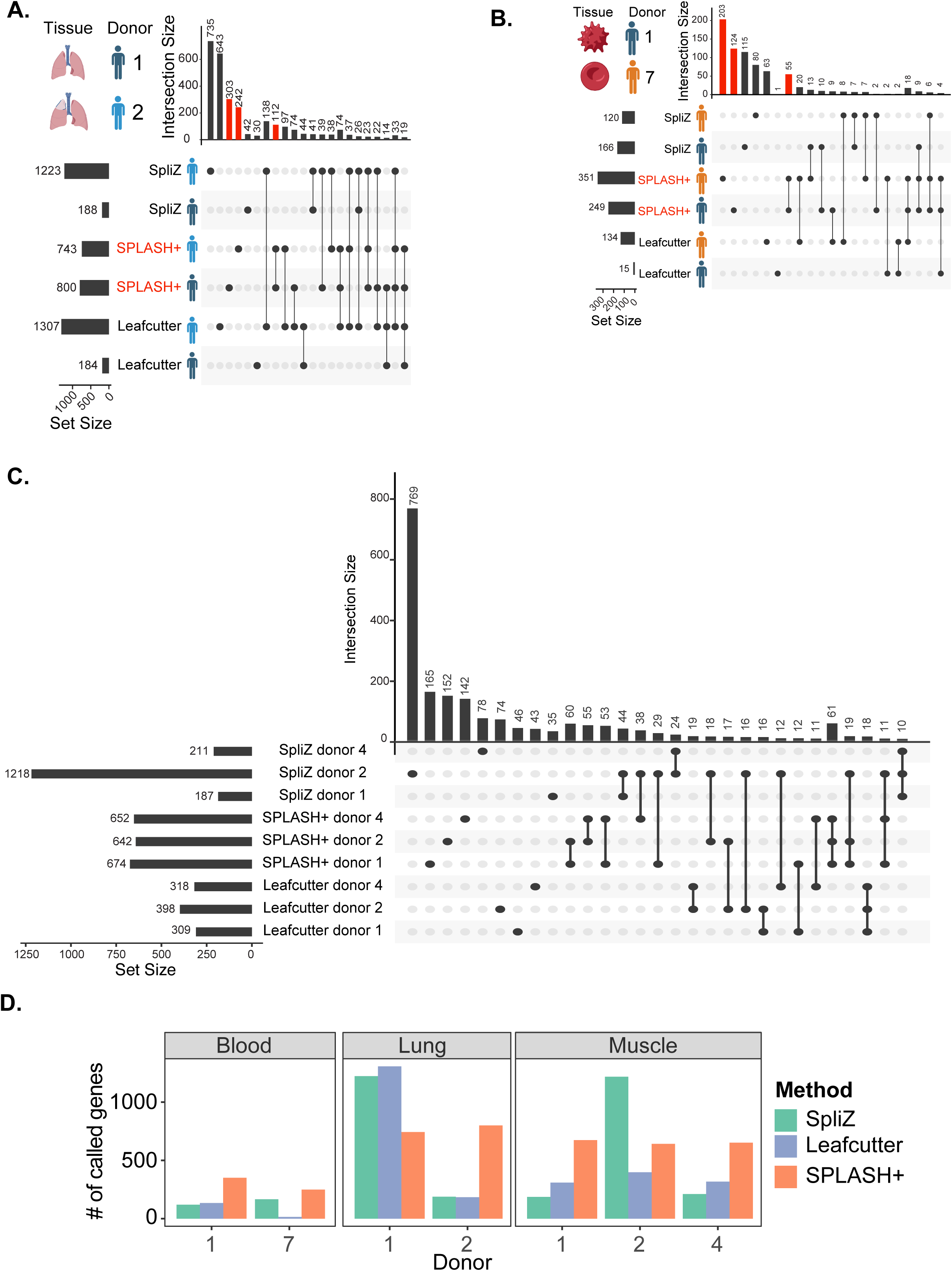
(A, B, C) Upset plots showing the comparison of the SpliZ, Leafcutter, and SPLASH+ for detecting genes with significant alternative splicing in lung, blood, and muscle tissues from each donor. (D) Barplots show the number of splicing genes called by each method and in each donor tissue.

**Suppl. Figure 10.**
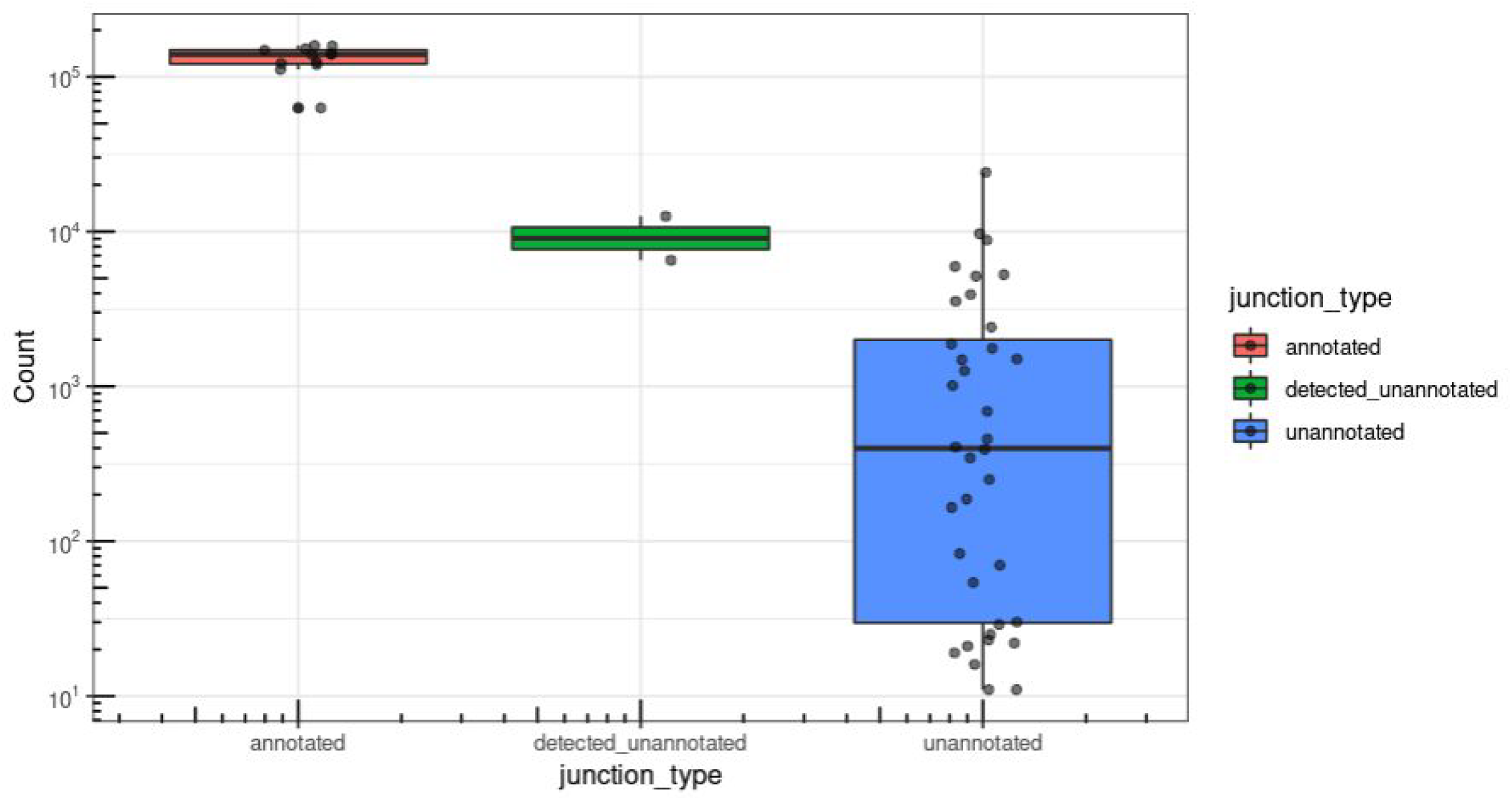
Number of SRA studies reported by Pebblescout for *CD47* junctions, providing further evidence for the expression of the two novel splice predictions made by SPLASH+ for *CD47*.

**Suppl. Figure 11.**
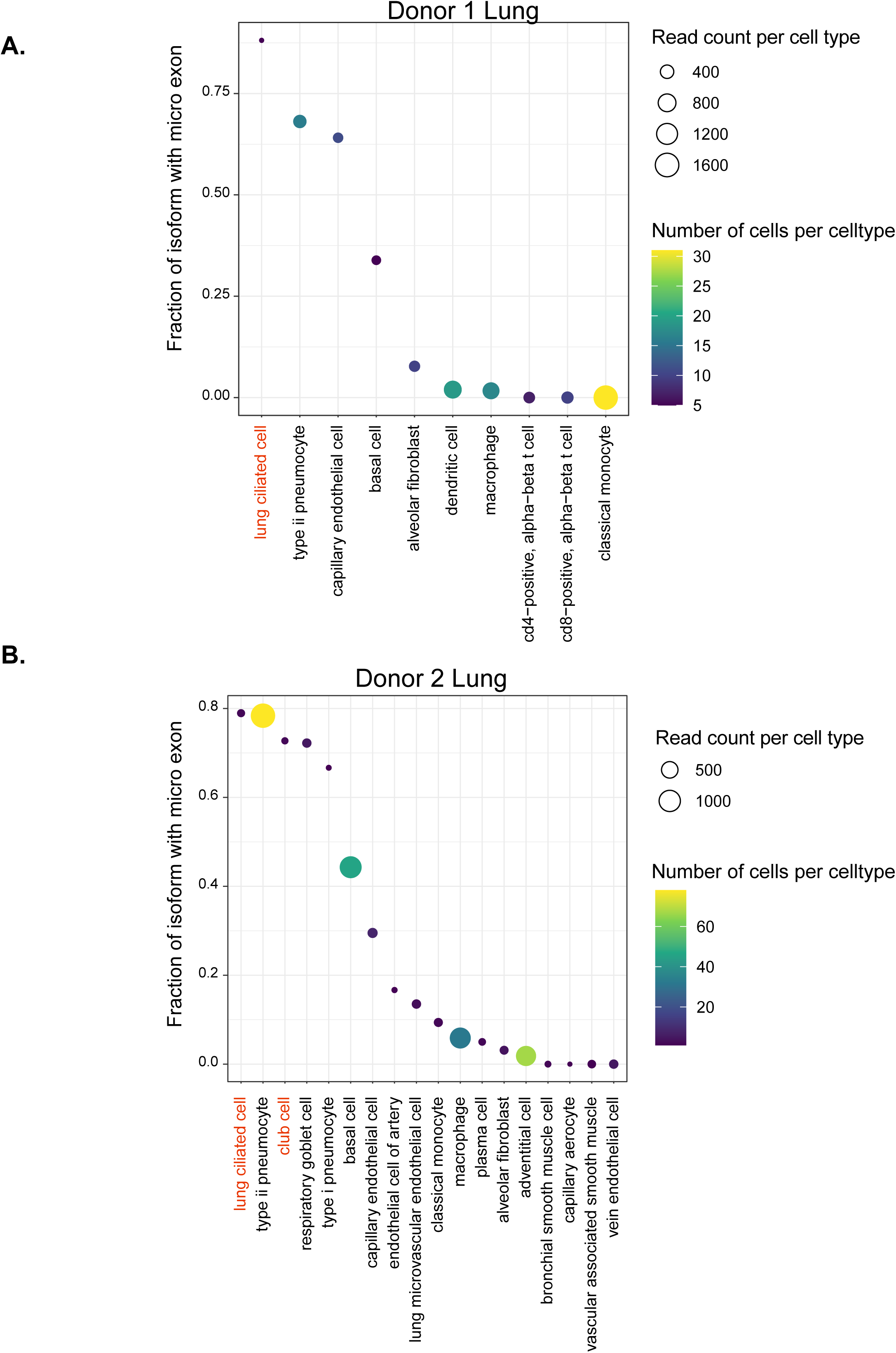
The two overrepresented cell types in bronchiole epithelium (ciliated and club cells) exhibit the highest expression fraction of the isoform with the microexon (dark blue isoform in Figure 3E) in lung replicates from donors 1 and 2, which is consistent with findings in the Visium data. Ciliated cells show 93 reads in 5 cells and 38 reads in 2 cells for donors 1 and 2, respectively, while club cells exhibit 22 reads in 2 cells for donor 2.

**Suppl. Figure 12.**
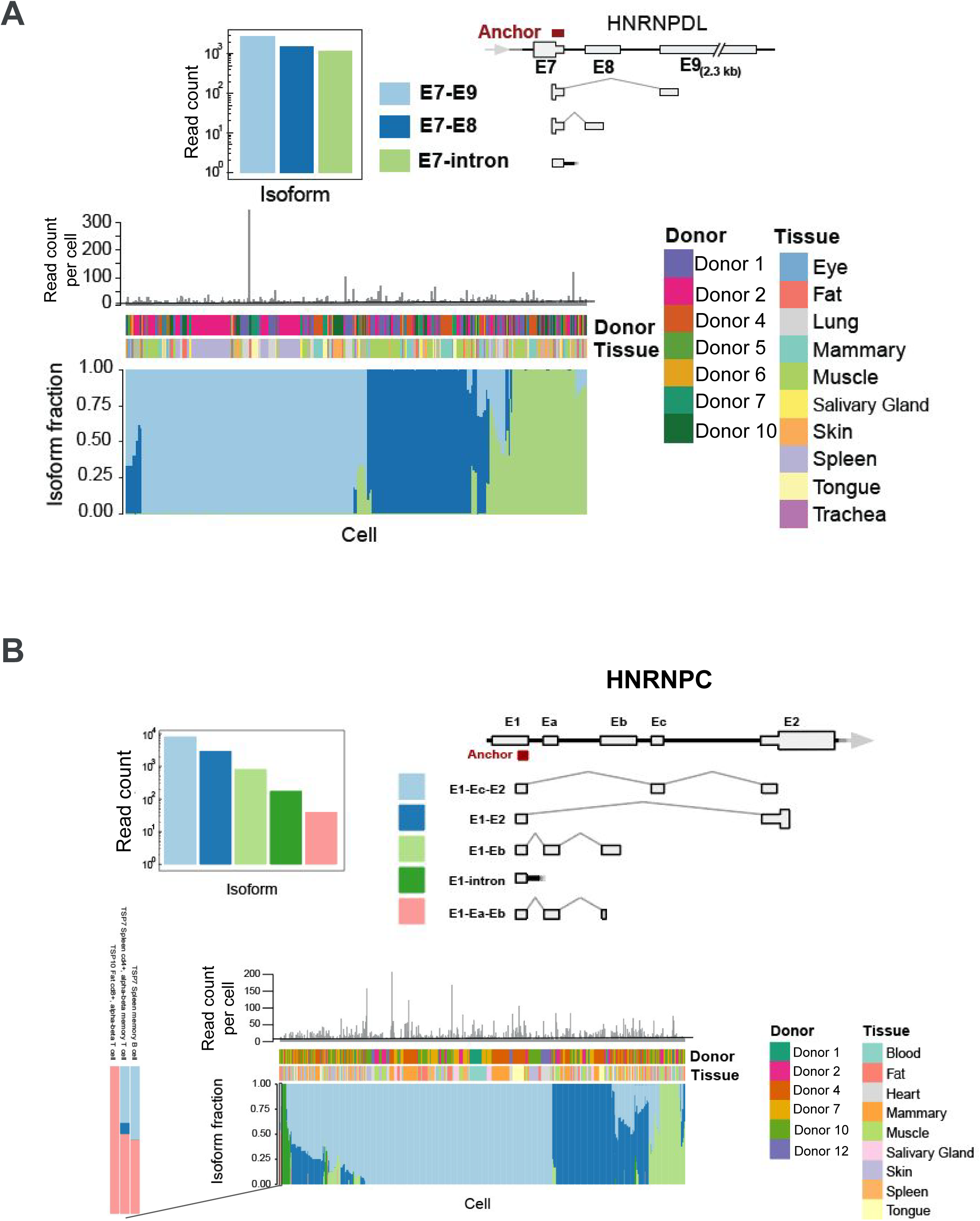
Single-cell dependent alternative splicing of two hnRNPs: (A) *HNRNPDL* and (B) *HNRNPC* illustrated for the anchor expressed in the most tissues for each of these genes.

**Suppl. Figure 13.**
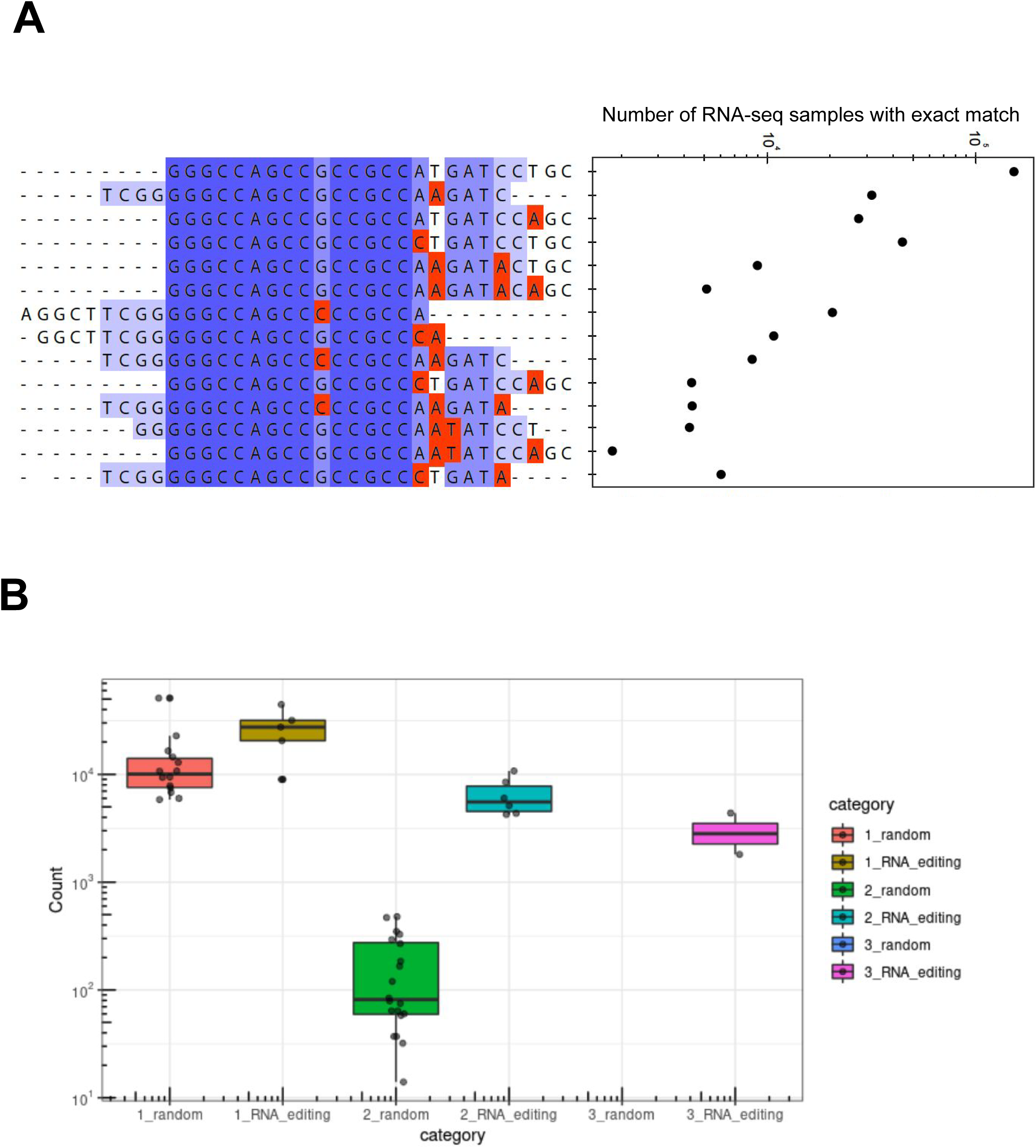
Query of the editing events detected in *ARPC2* using Pebblescout. (A) Number of studies reported by Pebblescout for the 25-mers with changes in the 6 RNA editing locations, as shown in Figure 6C. (B) Pebblescout reported compactors with editing in significantly more studies than decoy events or kmers constructed with base pair changes not predicted by SPLASH+.

## Tables

**Table 1: Compactor benchmarking.** This table contains the list of compactors that have matched to the fusions in simulated benchmarking datasets.

**Table 2: Centromeric anchors summary.** This table contains the sequence, read count, T2T CenSat annotation, and donor id, for each compactor of the centromeric anchors.

**Table 3: Noncoding RNA summary.** Number of called noncoding RNAs separated by each category for each donor and tissue.

**Table 4: Alternative splicing calls.** For each detected splicing anchor across the entire dataset, this table provides gene name, donor and tissue information, effect size, number of reads for the anchor, number of cells, the coordinates for the associated splice junctions, and whether the alternative splicing is annotated according to human T2T annotation.

**Table 5: SPLASH+ Comparison with SpliZ and Leafcutter.** This table contains significantly spliced genes detected by SPLASH+, SpliZ, and Leafcutter in lung, blood, and muscle tissues of *Tabula Sapiens* dataset.

**Table 6: Compactor diversity summary.** The number of distinct compactor sequences for each anchor in each donor.

**Table 7: SPLASH+ vs BraceR comparison.** This table contains the SPLASH+ IG compactor sequences along with their IgBLAST annotations and BraCeR contig sequences from the same cell for spleen and lymph node cells from donors 2 and 7 of *Tabula Sapiens*.

## METHODS

### Code availability

The code used in this work, including compactor generation, biological interpretation, is available at: https://github.com/salzman-lab/SPLASH-plus/. The code for splicing benchmarking can be found at: https://github.com/salzman-lab/SPLASH-plus/tree/main/Splicing_concordance_analysis

### Data availability

The FASTQ files for the Tabula Sapiens data were downloaded from https://tabula-sapiens-portal.ds.czbiohub.org/. Tabula Sapiens gene count table were obtained from the Tabula Sapiens AWS bucket. The Visium lung data was downloaded from SRA with accession ID SRR14851100.

Filtered BraCeR contigs for V(D)J analysis were downloaded from: s3://czb-tabula-sapiens/Pilot*/immune-repertoire-analysis/bracer. The five fusion simulated datasets for compactor benchmarking were downloaded from: https://data.broadinstitute.org/Trinity/CTAT_FUSIONTRANS_BENCHMARKING/on_simulated_data/sim_50/reads. The TPM values for simulated fusions were downloaded from: https://data.broadinstitute.org/Trinity/CTAT_FUSIONTRANS_BENCHMARKING/on_simulated_data/sim_50/metadata/sim_50.fusion_TPM_values.dat.

### SPLASH overview

SPLASH is a reference-free, annotation-free method that can be directly applied to raw sequencing reads and provide a unified statistical approach for the (co-)detection of various transcript diversification mechanisms (Chaung et al. 2023). Not requiring computational alignment of the reads to a reference genome, a feature commonplace in conventional RNA expression analysis methods, SPLASH can bypass inherent biases and blindspots in aligners. SPLASH takes sequencing files (FASTQ or FASTA) for input samples, and parses sequencing reads to find specific k-mers (substrings of length k) called *anchors* that are followed by a set diverse k-mers called *targets* across all reads from all input samples. In our analysis, we set the lengths for both anchors and targets to 27. SPLASH then builds a contingency table for each extracted anchor containing the read counts of each anchor’s target in each sample, i.e., each row and each column of an anchor’s contingency table corresponds to a target and a sample, respectively. Using these contingency tables, SPLASH then performs a statistical test for each anchor under the null hypothesis that the frequencies of anchor’s targets are independent of sample identity, yielding a closed-form valid p-value (Chaung et al. 2023). A significant p-value implies that the anchor has sample-dependent target distribution. The test statistic is constructed through random partitioning of the samples, and using random hash functions to map each target to a random value in [0,1]. As the test statistic is computed for different random choices of sample partitioning and random hashes per split, the p-value is Bonferroni-corrected to account for multiple testing across random sample partitioning and number of generated random hashes per split. SPLASH+ utilizes significant anchors for further downstream analysis by integrating a statistically valid local assembly approach, called compactors, and a framework for biological interpretation.

### SPLASH runs

Overall 19 tissues and 12 donors from the Tabula Sapiens dataset (Tabula Sapiens Consortium* et al. 2022) that have been profiled by SmartSeq2 were used for our analysis (Suppl. Figure 4). SPLASH was run in unsupervised mode on each donor and tissue separately. For donor tissues with >400 cells, we randomly selected 400 annotated cells to ensure implicit cell count normalization (and approximately read depth). Eight donor-tissues had fewer than 400 cells: donor 2 trachea (119 cells), donor 5 eye (134 cells), donor 1 blood (138 cells), donor 4 tongue (209 cells), donor 12 heart (277 cells), donor 3 eye (291 cells), donor 6 trachea (358 cells), donor 2 kidney (370 cells). In total, we ran SPLASH on 13,500 SmartSeq2 cells from 136 cell types. SPLASH was run with default parameters except for the number of random partitions of input cells (set to 300) and number of random hashes per partition (set to 10). Anchors with >50 reads in >10 cells that have SPLASH’s p-value < 0.05 and SPLASH’s effect_size > 0.2 were called as significant anchors.

### Compactor generation

To generate compactors for each anchor called as significant by SPLASH, first the input FASTQ files are searched for each anchor and the reads with exact match to the anchor are collected. The portion of the read that is upstream (left) of the anchor is clipped, i.e., for each anchor all the collected read sequences used for assembly start with the anchor sequence. Since there is no variation within the anchor, we always include the anchor as the starting sequence in the assembled sequence. Starting from the first position after the anchor, the frequency of each nucleotide is computed across the collected reads for the anchor and a new branch corresponding to a nucleotide whose frequency exceeds a certain threshold is created (Figure 1B). To create a new branch, we used the criterion that the nucleotide is in >10% of collected reads (if its count is >20) or in >80% of collected reads (if its count is >5). These thresholds are chosen based on the typical sequencing error rate and that we desire that the creation of a new branch due to sequencing error be highly unlikely, which is supported by our statistical analysis in the following section. Once a branch for a nucleotide is created at a given position, the subset of reads representing that nucleotide at that position are propagated to that branch (Figure 1B). If only one nucleotide satisfies the frequency criterion at a position, no extra branch is created and all reads are propagated to the subsequent iteration in the current branch. Corresponding to each position of a branch, the *compactor* sequence is defined as the anchor sequence extended by the nucleotides added at each position from the start of extension up to that position of the branch (Figure 1B). This process is repeated for each subsequent position and branch, using the reads propagated to each branch. It continues recursively until a user-specified number of recursions is reached (thereby fixing the compactor length for each set) or until the number of reads falls below a user-specified threshold. After this process, we report the set of compactors for each anchor where each compactor sequence corresponds to a specific path of branches and each compactor is reported with the count and set of reads representing it exactly.

### Compactor sequencing error model

Consider the compactor generation process, where we aim to decide on the nucleotide at a given position with N reads. Let *X_i_* ∈ {*A*,*T*,*C*,*G*}, 1 ≤ *i* ≤ *N* denote the nucleotide for read *i* at the given position. Under the null hypothesis, all variation in *X_i_* is due to sequencing error. Assuming that sequencing errors occur independently and uniformly across the sequenced regions at rate *∈*, considering without loss of generality that A is the ground truth nucleotide, we model the probability of observing each nucleotide in each read as:

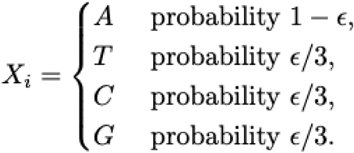

For each noisy nucleotide that is due to sequencing error (nucleotides other than A), the number of reads can be modeled as a binomial random variable with success probability 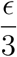 and *N* trials denoted by 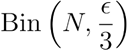. The probability of observing each noisy nucleotide {*T*,*C*,*G*} at least *d* times can be computed as the upper tail of 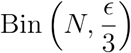, which can be upper bounded using Chernoff bound (Arratia and Gordon 1989) as follows:

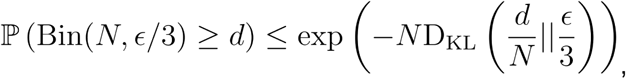

where D_KL_(*p*||*q*) denotes the Kullback–Leibler divergence between two independent bernoulli random variables with heads probabilities *p* and *q*:

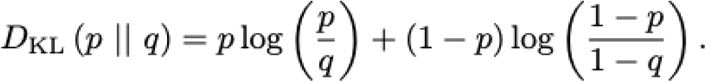

We can now compute a closed-form null probability for creating a new branch due to sequencing error at a given position with N reads and sequencing error rate *∈* when the criterion for generating a new branch for a nucleotide is to observe the nucleotide at that position in at least 10% of reads (i.e., 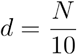). We use a conservative estimate of 1% for sequencing error rate. NovaSeq 6000 sequencing platform, the machine used to generate the Tabula Sapiens dataset, has a median error rate of 0.11% (Stoler and Nekrutenko 2021) and almost all studied datasets generated by NovaSeq 6000 had an error rate below 1% (Figure 2A in (Stoler and Nekrutenko 2021)). Under such sequencing error rate assumption, the null probability of creating a new branch for *N* = 50, 1000, 1000 is 0.012, 1.59×10^-4^, and 1.23×10^-38^, respectively, suggesting that it is highly unlikely to generate a new branch due to sequencing error when we consider a threshold of 10% for nucleotide frequency.

### Compactors benchmarking

We considered five simulated fusion datasets (sim_adipose, sim_brain, sim_colon, sim_heart, and sim_testis), each having 500 true positive (TP) fusions. We downloaded the ground truth sequences for TP fusions from: (https://data.broadinstitute.org/Trinity/CTAT_FUSIONTRANS_BENCHMARKING/on_simulated_data/simulated_fusion_transcript_sequences/). We used the breakpoint position provided for each TP fusion sequence in the downloaded files to obtain the seed anchor for each fusion, which was then used by compactors to reconstruct the fusion sequence. For each TP fusion, the seed anchor is extracted from its sequence as the sequence from `breakpoint_pos-26` to `breakpoint_pos` (Suppl. Figure 2A). We then employed these anchors and generated compactors by using the FASTQ files (we used R1 FASTQ files for compactor generation) (Suppl. Figure 2A). In order to find compactors that have evidence for TP fusion junctions, for each TP fusion we extract a junctional 20-mer sequence (10 bp on each side of the `breakpoint_pos` from the ground truth fusion sequence). Generated compactors that have a match to the 20-mer junctional sequences are then selected as the compactors with evidence for TP fusions. We report the number of fusions identified by compactors for each dataset as the number of unique fusions with at least one matched compactor. We discarded 185 fusions from our benchmarking analysis that did not have reads in R1 FASTQ files for their corresponding seed anchors.

### Classification of anchors into biologically relevant categories

To increase interpretability of the called anchors and to facilitate targeted downstream analysis for specific applications, we designed a biological interpretation step in SPLASH+ to assign a biologically relevant RNA diversifying event to each anchor based on the features directly derived from its compactors. We consider six categories for anchors (Suppl. Figure 3): splicing, internal splicing, base pair change, 3’UTR, centromere, and repeat. If an anchor is not assigned to any of these categories, it will be categorized as an unclassified anchor. We used a hybrid approach to assign classes to anchors, with some classes assigned independently of reference genome alignment (e.g., internal splicing and base pair change) and others assigned based on alignment (e.g., splicing, 3’UTR, centromere, and repeat). As each anchor might be qualified for more than one class, we prioritize classes in the following order: splicing, internal splicing, base pair change, 3’UTR, centromere, repeat. To classify anchors, we consider only the top two most abundant compactors for each anchor (i.e., those with the highest fraction of anchor reads). If one compactor is longer than the other, we consider its substring equal in length to the shorter compactor. We then compute two different string metrics: Hamming distance and Levenstein distance. We should note that both Levenstein and Hamming distance for the two compactors are computed after removing the anchor sequence. Anchors with the same Hamming and Levenstein distances are classified as base pair change as this criterion suggests that only substitutions (i.e., nucleotide changes) account for the difference between the two compactors. We also utilize the sequence of Levenstein operations (comprising insertions, deletions, and substitutions) and classify an anchor as “internal splicing” if the Levenstein distance is less than the sum of of the longest stretch of deletions (*Run_D*) and insertions (*Run_I*) plus one (Suppl. Figure 3).

Other anchor categories (such as splicing) are assigned based on the alignment of compactors to the reference genome. To identify anchors potentially explained by alternative splicing, we align the two compactors for each anchor to the T2T human reference genome using STAR v2.7.5.a (Dobin et al. 2013) with the following parameters:

`--twopassMode Basic --alignIntronMa× 1000000 --chimJunctionOverhangMin 10`
`--chimSegmentReadGapMax 0 --chimOutJunctionFormat 1 --chimSegmentMin 12`
`--chimScoreJunctionNonGTAG −4 --chimNonchimScoreDropMin 10 --outSAMtype SAM`
`--chimOutType SeparateSAMold --outSAMunmapped None --clip3pAdapterSeq AAAAAAAAA`
`--outSAMattributes NH HI AS nM NM`

We then extract information about the mapping flag, chromosome, coordinate, CIGAR string, and number of mismatches from the STAR BAM file (1st, 2nd, 3rd, 4th, 6th, and 16th columns). If at least one of the two compactors involves a split alignment, and the Hamming distance and Levenstein distance are not equal (as this would indicate a base pair change), we classify the anchor as “splicing”. This classification is based on the presence of a splice junction in at least one compactor and that the difference between the compactor sequences cannot be explained by simple substitutions. Note that both compactors should overlap with the same gene to be considered as splicing anchor.

Anchors not classified as splicing, base pair change, or internal splicing are considered for further classification. We intersect compactor mapping positions with annotated 3’UTRs, centromere satellite elements (from T2T CenSat database), and repetitive elements (from RepeatMasker database) using BEDTools v2.25.0 (Quinlan 2014) intersect command. Anchors are then categorized to one of these categories based on this intersection (Suppl. Figure 3).

### Comparison to SpliZ and Leafcutter for alternative splicing

SpliZ is a statistical method for detecting genes with cell type-specific alternative splicing in scRNA-Seq (Olivieri, Dehghannasiri, and Salzman 2022). It assigns a single score to each pair of cell and gene and is reference-dependent in the sense that it needs the split reads mapping to the splice junctions of the gene. To obtain high confidence splice junctions needed for SpliZ analysis, we first aligned reads to human reference genome using STAR and then ran SICILIAN (Dehghannasiri, Olivieri, and Salzman 2020), a statistical wrapper for detecting high-confidence splice junctions from spliced aligners, on the STAR BAM files. We applied SpliZ to the reads aligned to the detected splice junctions. SpliZ was run on each donor separately and to avoid calling genes with tissue-specific splicing rather than cell type-specific splicing, its statistical test was performed separately across cell types within each tissue from that donor. Both SpliZ and SICILIAN were run with default parameters.

For performing splicing analysis using Leafcutter (Li et al. 2018), we first extracted junctional reads by running RegTools v0.5.2 (Cotto et al. 2023) on STAR BAM files with parameters suggested on Leafcutter GitHub (https://davidaknowles.github.io/leafcutter/articles/Usage.html): minimum anchor length of 8bp on each side of the junction, minimum intron length of 50 and maximum intron length of 500 kb. Junctions from each cell within a donor tissue were then clustered (i.e., clusters with overlapping exon start/ends) using Leafcutter clustering script (`leafcutter_cluster_regtools.py`) with default value of 50 minimum total reads per cluster to create a matrix of junction counts across all cells within each donor tissue which are subsequently used for differential splicing analysis. Since Leafcutter is a supervised method for differential analysis between samples from two groups, we performed differential splicing analysis for each pair of cell types within a donor tissue that had at least five cells. We chose this threshold because Leafcutter warns that p-values are not calibrated for groups with fewer than four samples: “p-values are not calibrated for less than four samples” (though it is still possible to run on less than four samples)”. To annotate intron clusters called by Leafcutter, we used the `gtf_to_exons.R` script (downloaded from Leafcutter repository) on the T2T annotation GTF file to create an exons file required for Leafcutter annotation. Intron clusters with an adjusted p-value (p.adjust) < 0.05 and an absolute log effect size (logef) > 1.5 were considered significant. A gene is called as significantly differentially spliced in a donor tissue by Leafcutter, if it had at least one significant intron cluster in any tested cell type pair for that donor tissue.

We tested the significance of the overlap in splicing genes called by SPLASH+ and each of the other two methods (Leafcutter and SpliZ) for every donor tissue. Let *N_1_* and *N_2_* be the number of splicing genes called by SPLASH+ and the other method, respectively in a donor tissue. Under the null hypothesis, the probability that a gene is identified by both methods is 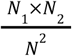, where *N* is the total number of genes. Observing *N_s_* genes identified by both methods, we can now compute the binomial p-values to test the significance of this overlap. We report the -log10(p-value) for the significance of the overlap between SPLASH+ and SPliZ, and between SPLASH+ and Leafcutter for each donor tissue (Figure 3B, bottom). The resulting p-values were significant for all comparisons, with the overlap between SPLASH+ and Leafcutter showing greater significance than the overlap between SPLASH+ and SpliZ for all donor tissues except for blood donor 1.

### Querying SRA human RNA-Seq database using Pebblescout for novel splicing and RNA editing

Pebblescout (Shiryev and Agarwala 2024) is a kmer-based query tool designed to take input sequences and report studies (RNA-Seq sequencing samples) within its indexed database that share matches with the provided input. Pebblescout randomly samples a 25-mer from every 42-mer in the query sequence, guaranteeing a match between the query and the study if the match is at least 42bp long. Given that Pebblescout reports matches based on any 25-mer in the input sequence, for each queried *CD47* and *HSP90AA1* isoform a 42-mer (the minimum acceptable length by Pebblescout for a query) is constructed by taking 21 bps from both the 5’ and 3’ sides of the junctions. For queries regarding *ARPC2* (Figure 6C), we extended each 20-mer by 22 bps. We further aligned the reported 25-mer matches by Pebblescout to the reference genome using Bowtie2 v2.2.1 to confirm their unique mapping to the *CD47*, *HSP90AA1*, and *ARPC2* loci.

To create a control set of unannotated junctions for novel *CD47* and *HSP90AA1* junctions, we first extracted exonic 21 bps leading to each annotated 5’ and 3’ exon boundary for each gene. We then concatenated the 21 bps from each 5’ splice site with those from its downstream 3’ splice sites to form 42-mers. These junctions that are not part of the annotated gene structure were then used as a control set for evaluating the prevalence of detected unannotated junctions by SPLASH+.

### V(D)J rearrangement analysis and comparison to BraCeR

We defined immunoglobulin compactors (being referred to as “IG-compactors”) as the compactors mapped to an immunoglobulin gene *IGH*, *IGK*, or *IGV* by STAR. We further defined immunoglobulin-anchors (“IG-anchors”) as anchors with >20% of reads associated with IG-compactors. The unaligned compactors of an IG-anchor are still considered IG-compactors through “annotation-by-association”. To compare the SPLASH+ with BraCeR (Lindeman et al. 2018), we considered a stringent criterion for IG-compactors where we first annotated IG-compactors with AssignGenes.py (ChangeO v1.3.0) (Gupta et al. 2015) and IgBLAST v1.21.0 (Ye et al. 2013) and considered only those IG-compactors that had both variable (V) and joining (J) immunoglobulin gene segments identified through IgBLAST. The IG-compactors are annotated as in-frame by IgBLAST if the last triplet of the annotated V gene is in-frame with the first triplet of the annotated J gene. Also, to test whether SPLASH+ provides support for the detection of the same B cell receptors (BCRs) as BraCeR, for each cell we computed the minimum Hamming distance between the compactors of IG-anchors and the filtered BraCeR contigs.

### Repeat expansion analysis

For each cell, we obtain total read count for each repeat length by summing the reads across all compactors associated with that repeat length. To characterize the repeat length for the two dominant alleles per cell, we refer to the repeat length with the highest read count as the “primary repeat length” and the repeat length with the second-highest read count as the “secondary repeat length”.

### Analyzing the abundance of the third most abundant target under a two-true-target assumption

We statistically quantify how unlikely it is that we observe a high count for the third most common target when there are only two ground truth targets and all others are due to sequencing error. This could arise when the anchor corresponds to alternative splicing involving two isoforms (each target represents an isoform) or corresponds to an SNP (one target represents the transcript without SNP and the other target represents the transcript with SNP). We assume that the anchors are observed without sequencing error. We also make the same assumption of uniformity and independence for sequencing error rate *∈* as discussed in the section on analyzing sequencing errors in compactor construction. Under these assumptions, we can provide p-values on the abundance of the third most frequent target, showing that it is very unlikely to observe a high count for the third most abundant target when only two ground truth targets exist, with all others attributed to sequencing errors.

Let *M* be the total number of counts, and *L* be the target length. The two ground truth targets are denoted by *y*_1_ and *y*_2_. For each possible target (a sequence of length *L* from A/G/C/T) *y* ∈ {*A*, *C*, *T*, *G*}*^L^*, we define *l*(*y*) to be the minimum of the hamming distance between and *y*_1_, and *y* and *y*_2_, i.e., *l*(*y*) = min (*d*_H_(*y*,*y*_1_), *d*_H_(*y*,*y*_2_)), where *d*_H_ denotes the hamming distance between two strings. Note that *l*(*y*) is always between 1 and *L*. We can now compute the probability of observing a noisy target *p_y_* due to sequencing error using its minimum hamming distance *l*(*y*) relative to the two true targets. For this target *y*, assuming that targets are generated randomly from *y*_1_, *y*_2_ and that ℙ(unerrored target *y*_1_) = *q*:

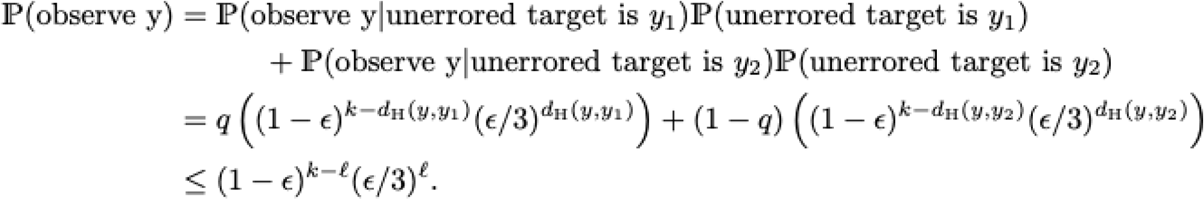

We used the law of total probability for the first equality, and the last upper bound is because both hamming distances *d_H_*(·) are greater than *l*.

We can now compute probability of observing more than *k* counts for the third most abundant target. Let *X*_1_, *X*_2_, …, *X*_4_^*L*^ to be the counts of the 4*^*L*^* possible targets. We consider *X*_1_ and *X*_2_ to be the counts for the top two true targets *y*_1_ and *y*_2_, respectively. Under the null, *X*_3_, …, *X_k_* are due to sequencing error, and we assume these are arbitrarily ordered (e.g., lexicographically), with the target sequence corresponding to *X_i_* being *y_i_*. Therefore, the counts for the third target is obtained as max(*X*_3_, …, *X*_4_*^L^*). We can now compute the probability of at least *K* counts for the third target as follows:

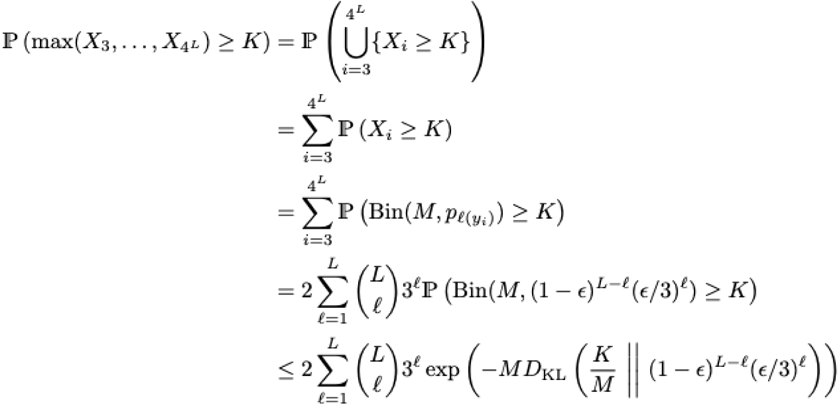

The first equality implies that in order for the maximum count of noisy targets be greater than *K*, at least one noisy target must have counts greater than *K*. The third equality states that *X_i_* can be modeled as a Binomial random variable 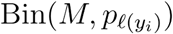 using the total count and probability of observing *y_i_* which is a function of its minimum hamming distance to the top two targets. The fourth equality is because there are 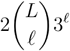 targets with minimum hamming distance *l*. For *y*_1_, there are exactly 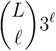 targets of hamming distance exactly *l* from *y*_1_ (choose *l* out of the *L* possible locations, and which of the 3 nucleotides to observe at each of the *l* locations), and therefore there are at most 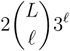 targets with minimum hamming distance *l* between *y*_1_, *y*_2_. The final inequality utilizes a Chernoff bound (Arratia and Gordon 1989) for the binomial distribution. This bound we obtained for the count of the third target *K* decays very quickly as a function of *K*. For example, for *M* = 50 and *∈* = 0.01, *K* = 4 yields .01, *K* = 5 yields 1*E* - 5, and *K*= 7 yields 1*E* - 10. We have provided a rigorous statistical framework to show that it is very unlikely to observe many counts for noisy targets that are due to sequencing error.

## Notes

### Competing Interest Statement

The authors have declared no competing interest.

### Summary of Updates

In this version, we have made multiple revisions to further enhance results and figures in the preprint.

